# Epigenetic changes and serotype-specific interferon-responses of lung epithelial cells in late post-influenza pneumococcal pneumonia

**DOI:** 10.1101/2023.06.28.546771

**Authors:** Julia D Boehme, Andreas Jeron, Kristin Schultz, Lars Melcher, Katharina Schott, Elif Gelmez, Andrea Kröger, Sabine Stegemann-Koniszewski, Dunja Bruder

## Abstract

Pneumococcal infection following influenza A virus (IAV) pneumonia is a synergistic complication with high mortality. IAV modulates host antibacterial responses and invasiveness of pneumococcal serotypes and is an important pathogenic factor ^2^. Yet, serotype-specifc immediate-early responses of the IAV-perturbed alveolar epithelium have not been adressed. We analyzed gene transcription in alveolar type II epithelial cells (AECII) from mice infected with IAV and/or one of three *S. pneumoniae* (*S.pn.*) serotypes of varying invasiveness (4 > 7F > 19F). IAV, 14 days post infection, rendered the lung susceptible to invasive *S.pn.* infection with serotype 4 and the mildly invasive 7F but not 19F. Only 7F secondary infection induced exacerbated cytokine/chemokine responses. IAV/7F infection induced superior protein expression of type I and II interferons, exceeding that in IAV/serotype 4 infection. Inference of a scale-free-like ARACNE gene co-expression network revealed interferon-response network modules. Network-mapping unfolded *S.pn.* serotype-specific transcriptional network responses/usage. Secondary *S.pn.* infection abrogated the IAV-induced pneumocyte proliferative configuration and IAV infection rendered the transcriptional response to 7F comparable to that of serotype 4. This related to network genes correlating with the expression of two master regulators of interferon responses: *Irf7* and *Stat1*. Epigenetic ATAC-seq analysis of AECII in resolved IAV infection identified enhanced expression of ARACNE network genes *Hist1h2bf*, *Igtp*, *Mki67*, *Rasl10b*, *H2-Q6* and *H2-Q7* to be associated with increased chromatin accessability at promoter regions. We show that AECII retain a sustained IAV-associated transcriptional configuration with epigenetic involvement that serotype-specifically affects proliferation and intensifies the AECII transcriptional response, mainly to interferons, in *S.pn.* infection.

## Introduction

Influenza (flu) viruses are respiratory pathogens with high zoonotic potential estimated to cause over 5 million hospitalizations worldwide per year ^1^. Despite the development and annual adjustment of influenza vaccines, high incidences of flu disease are recorded during seasonal epidemics as well as during the most recent swine flu pandemic in 2009 ^2^. Influenza A virus (IAV) and influenza B virus are the most common flu subtypes causing influenza-related disease in humans ^3^.

Importantly, bacterial secondary infections are frequent complications of influenza disease and are associated with high morbidity and mortality^3^. *Streptococcus pneumoniae* (*S.pn.*) is a Gram-positive bacterial pathogen and a frequent, mostly asymptomatic colonizer of the human nasopharynx. Epidemiological evidence attested *S.pn.* to be one of the major pathogens identified in post-influenza secondary bacterial infections ^4, 5^. Experimental data revealed that a variety of pathogen- and host-specific mechanisms, *e.g.* increased attachment ^6^, availability of host-derived nutrients ^7, 8^ or host antiviral immune responses ^9, 10^ critically drive influenza/pneumococcal co-pathogenesis. Despite intense research over the last two decades, the detailed mechanisms of this co-pathogenesis remain elusive in many points. *S.pn.* comprises over 94 serotypes, defined by the organo-chemical structure of their polysaccharide capsule. *S.pn.* in its own right causes invasive pneumococcal disease (IPD), a severe respiratory bacterial infection manifesting *e.g.* as pneumonia, otitis media or meningitis. In 2018, the European Centre for Disease Prevention and Control (ECDC) reported 24,663 confirmed cases of IPD mainly caused by serotypes 8, 3, 19A, 22F, 12F, 9N, 15A, 10A, 23B and 6C ^11^. Most *S.pn.* serotypes colonize the nasopharynx asymptomatically and are frequently isolated therein. Generally, invasive serotypes are found less frequently in the nasopharynx of healthy individuals. Mechanistic understanding of the colonizing/invasive potential of *S.pn.* serotypes is however largely missing.

Invasive serotypes of *S.pn.* were shown to produce thinner, energetically more costly capsules as compared to colonizing serotypes^12^. Interestingly, by exchanging serotype-specific capsule operons, the according serotype’s growth phenotype in nutrient-restricted conditions is also transferrable ^12^. Invasiveness of *S.pn.* serotypes also depends partly on the resistance to complement factor 3 (C3) deposition onto the bacterial surface, inducing phagocytosis and killing by innate immune cells ^13^. In this regard, the serotype-defining capsule type is more relevant than other genetic variations between *S.pn.* serotypes ^13^.

Available *S.pn.* vaccines immunize against various serotypes. The currently used 23-valent Pneumovax vaccine contains antigenic determinants of serotypes 1, 2, 3, 4, 5, 6B, 7F, 8, 9N, 9V, 10A, 11A, 12F, 14, 15B, 17F, 18C, 19F, 19A, 20, 22F, 23F and 33F ^14^. In humans, serotypes 1, 4, 5, 7F, 8, 12F, 14, 18C, and 19A are considered to have high invasive and bacteremic potential ^15^.

The vast majority of *in vivo* studies addressing the lethal synergism between influenza viruses and *S.pn.* employ infection models, in which secondary pneumococcal infection is performed during acute viral pneumonia. In this phase of peak susceptibility, pulmonary integrity is greatly compromised by viral replication as well as antiviral immune responses.

However, we and others have previously shown that influenza-mediated susceptibility to *S.pn.* can persist after viral clearance has been accomplished, *i.e*. during the post-influenza recovery period ^16, 17^. In contrast to equal hypersusceptibility to both invasive and colonizing strains during acute influenza infection, we found persisting susceptibility to *S.pn.* only for invasive isolates during the post-flu recovery phase. Physiologically, this phase is characterized by gradual contraction of antiviral immune effectors including lung innate and adaptive leukocyte subsets and airway inflammatory cytokines (own unpublished data) and proliferation of alveolar type II epithelial cells (AECII) ^18^.

In the lower airways, AECII together with AECI comprise the epithelial barrier, with a subset of AECII acting as progenitor cells ^19^. With respect to IAV, AECII are primary sites of viral replication in the alveoli ^20^. They substantially shape immune responses against influenza and other respiratory viruses including poxvirus ^21^ and SARS-CoV2 ^22^ upon recognition of viral pathogen-associated molecular patterns (PAMPs) ^23^ as well as antiviral alarmins such as type I interferons (IFNs). Thereby, AECII, next to other long-lived locally residing cells such as macrophages and dendritic cells (DC), contribute to the sentinel system of the lower airways. Currently, little is known about AECII-specific processes in the post-IAV recovery phase and how they may interfere with immediate-early recognition and defense responses to secondary bacterial invadors such as *S.pn*.

Inflammatory priming and trained immunity describe mechanisms of sustained imprinting of secondary immune responses by primary triggers. These can be mediated by epigenetic, transcriptional and metabolic changes to resident (immune) cells and their progenitors altering their immunological responsiveness ^24^. To date, it remains unclear whether and how such mechanisms contribute to sustained enhanced susceptibility to *S.pn.* following influenza infections, especially with regard to structural cells such as AECII.

Here, we employed a murine model of post-influenza secondary pneumococcal infection to analyze *S. pn.* serotype-specific AECII-responses in the regenerating lung. Thereby we aimed at determining the contribution of influenza-experienced epithelial cells to aberrant antimicrobial immune respones during secondary pneumococcal pneumonia. We confirmed that resolved IAV infection enhances susceptibility to pneumococci in a strain-dependent manner, partly associated with the emergence of a serotype-specific hyperinflammatory airway microenvironment. On the AECII transcriptional level, we found that abrogation of post-flu AECII epithelial repair was a general consequence of pneumococcal secondary infection. AECII from IAV-affected lungs furthermore mounted faster and intensified responses to type I and II interferons induced by the secondary *S.pn.* infection and showed ATAC-based epigenetic adaptations of their transcriptional response.

## Materials and Methods

### Mice

For all experiments, female C57BL/6JOlaHsd mice (age 10 - 14 weeks) from Envigo were used. All mice were housed in the animal facility of the Helmholtz Centre for Infection Research under specific-pathogen-free (SPF) conditions and in accordance with national and institutional guidelines. The *in vivo* study protocol was reviewed and approved by institutional and regional ethical bodies (Niedersaechsissches Landesamt fuer Verbaucherschutz und Lebensmittelsicherheit).

### Influenza infections

Madin-Darby canine kidney cell-derived IAV (H1N1, PR/8/34, mouse-adapted) was obtained^25^ and the 50 % tissue culture infectious dose (TCID_50_) was determined as previously described ^26^. Mice were anesthetized by intraperitoneal (i.p.) injection of ketamine/xylazine and were intranasally infected with 7.9 TCID_50_ in 25 µL of PBS. Control animals received PBS only.

### Pneumococcal infection

For *S. pneumoniae* infections, the serotype 4 strain TIGR4 (ATCC BAA-334) ^27^, a serotype 19F strain (BHN100) ^28^ and a serotype 7F strain (BHN54) ^28^ were used. Strains were obtained from B. Henriques-Normark (Karolinska Institutet, Stockholm, Sweden). Bacteria were plated from frozen glycerol stocks onto Columbia blood agar plates (BD) and incubated for 12 - 18 h (37 °C, 5 % CO_2_). Single colonies were resuspended in fresh Todd-Hewitt yeast medium and bacterial suspension was grown to mid-logarithmic phase (OD600 nm ∼ 0.4), bacteria were spun down, washed once and diluted in PBS to obtain the final inoculum for infection. CFU were retrospectively determined by plating. For bacterial infections, mice were anesthetized by i.p. injection of ketamine / xylazine and 40 µL of the pneumococcal inoculum (or PBS) were administered by oropharyngeal aspiration.

### AECII isolation

Isolation of primary murine AECII was conducted according to a previously established protocol ^29^ with slight modifications. Briefly, lung single cells suspensions were obtained by enzymatic (dispase / DNase) and mechanical dissociation of whole lungs. Pooled cells from n = 3 - 5 mice/cohort were stained with PE- or APC-conjugated antibodies (from either BD or BioLegend) against murine F4/80 (clone: BM8), CD93 (clone: AA4-1), CD11c (clone: N418), CD19 (clone: 6D5), CD31 (clone: 390), CD11b (M1/70), CD16/32 (clone: 2.4G2) and CD45 (clone: 30-F11). AECII were sort-purified by negative selection (PE^-^, APC^-^, autofluorescence^hi^, side-scatter^hi^) using FACS Aria II and Fusion instruments (BD) and maximum purity masks (4-way purity mode).

### Generation of bronchoalveolar lavage fluid (BALF) samples

At experimental endpoints mice were euthanized, the trachea was exposed, and an indwelling venous catheter was inserted. The lungs were flushed with 1 mL ice-cold PBS via the catheter using a syringe. Where appropriate, obtained BALF was used for CFU quantification, then spun down and BALF supernatants were stored at -80 °C for further analyses.

### CFU quantification

At experimental endpoints serial dilutions of BALF and blood samples of *S.pn.* infected mice (with and without prior IAV infection) were plated onto Columbia blood agar plates (BD) to assess airway bacterial burden and bacteremia.

### Albumin, cytokine and chemokine quantification

Airway serum albumin was quantified by ELISA using BALF samples and anti-mouse albumin coating- and HRP-conjugated anti-mouse albumin detection antibodies (both Bethyl Laboratories) and a purified mouse serum albumin standard (Sigma Aldrich). Airway cytokines and chemokines were quantified by multiplex analyses of BALF samples using LEGENDplex^TM^ kits (BioLegend). Quantification of airway bioactive type I/III IFN was conducted by treating type I/III IFN-sensitive cells isolated from Mx2-Luc reporter mice for 24 h with diluted BALF. Relative IFN levels were determined by quantifying relative light units (RLU) in cell lysates after addition of luciferin-reaction buffer ^30^. IFN-α was quantified using the VeriKine-HS^TM^ Mouse Interferon Alpha All Subtype ELISA Kit (PBL Assay Science^TM^). IFN-β and IFN-λ2/3 were quantified using the respective Mouse DuoSet ELISA kits (R&D Systems^TM^).

### Western blot analyses

FACS-sorted, pooled AECII were lysed in SDS sample buffer (4 % SDS, 1.5 M TrisHcl, 33 % Glycine, 6 % β-Mercaptoethanol, 0.5 % bromophenol blue) at 95 °C for 3 - 5 min. Cell lysates were loaded on 10 % acrylamide gels, proteins were separated and blotted on PVDF membranes. The following RRID-listed primary antibodies were used for detection: anti-IRF1 (AB_631838), anti-IRF3 (AB_2264929), anti-IRF7 (AB_1125072), anti-IRF9 (AB_2296227), anti-IL10RB (AB_1512256), anti-IFNGR2 (AB_2248682), anti-STAT1 (AB_2198300), anti-STAT2 (AB_2799824), anti-vinculin (AB_2819348) and anti-GAPDH (AB_2107448).HRP-conjugated secondary antibodies reactive against either mouse, rabbit, goat or hamster Ig were utilized. Blots were developed using Lumi-Light western blotting substrate (Millipore) and imaged using the INTAS ECL Chemocam Imager (Intas Science Imaging). Average band intensities were normalized against the respective loading controls using ImageJ software.

### Transcription analysis

Total RNA from FACS-sorted, pooled AECII (n = 3 independent replicates per experimental condition, cells pooled from n = 3 - 5 mice / cohort) was isolated using the RNeasy Plus Mini Kit (Qiagen). Clariom S microarray analysis (Affymetrix, Thermo Fisher Scientific) was performed at the Helmholtz Center for Infection Research (Braunschweig, Germany) according to manufacturer’s recommendations. Raw data were analyzed with the Transcriptome Analysis Console (Thermo Fisher Scientific) using the SST-RMA algorithm. Transcripts with log_2_ signal intensities (SI) below the 20^th^ percentile of the overall SI-distribution in all 42 microarrays were excluded from further analysis. Differential gene expression was calculated versus the mean of the PBS control replicates. Transcripts with a fold change (FC) of |FC| > 3 and ANOVA-based p-value of p < 0.05 were considered significant. Normalized log_2_ SI data of these transcripts were further processed to infer a gene co-expression network based on the ARACNE algorithm ^31^, implemented in the Cyni Toolbox addon ^32^ of the Cytoscape analysis software ^33^. The ARACNE algorithm settings were as follows: Mutual information (MI) calculation was performed using the adaptive partitioning option^34^. DPI tolerance was set to 0. As a regulatory basis for the ARACNE network, a list of all known murine transcription factors was used, that was compiled from online resources from the Gene Ontology Consortium. The MI-threshold was empirically set to 0.6, resulting in a network with 449 nodes and 791 edges with three network components. Minor unconnected network components as well as unconnected individual nodes were excluded from further analyses. The resulting ARACNE gene co-expression network was further analyzed using Gephi software ^35^ for node arrangement by multi-gravity force atlas algorithm ^36^, modularity/spectral network partitioning ^37^ and calculation of basic network characteristics like node connectivity, clustering coefficient and node betweenness centrality. For the visualization and plotting of network figures and associated characteristics a series of Python scripts were used. For ANOVA analysis, visualization and clustering of gene expression and gene ontology results, Genesis software ^38^ was used. Gene ontology analysis was performed using the ClueGO plugin for Cytoscape Software. For receptor/ligand mapping, the database from CellTalkDB ^39^ together with a Python script was used. Microarray data were deposited online and are available at NCBI’s Gene Expression Omnibus (GEO) under reference ID: GSE225343. Python scripts for the analysis and visualization of transcriptomics data and network representation are available on reasonable request.

### ATAC-seq analysis

FACS-sorted AECII (pooled from 5 - 6 mice/experimental group) were frozen in culture medium containing FBS and 5 % v/v DMSO. Cryopreserved cells were sent to Active Motif Inc. for ATAC-sequencing. Cells were thawed in a 37 °C water bath, pelleted, washed with cold PBS, and tagmented as previously described ^40^, with some modifications based on Corces *et al.* (2017) ^41^. Briefly, cell pellets were resuspended in lysis buffer, pelleted, and tagmented using the enzyme and buffer provided in the Nextera Library Prep Kit (Illumina). Tagmented DNA was purified using the MinElute PCR purification kit (Qiagen), amplified with 10 cycles of PCR, and purified using Agencourt AMPure SPRI beads (Beckman Coulter). Resulting material was quantified using the KAPA Library Quantification Kit for Illumina platforms (KAPA Biosystems) and sequenced with PE42 sequencing on the NextSeq 500 sequencer (Illumina). Reads were aligned using the BWA algorithm ^42^ (mem mode; default settings). Duplicate reads were removed, only reads mapping as matched pairs and only uniquely mapped reads (mapping quality ≥ 1) were used for further analysis. Alignments were extended *in silico* at their 3’-ends to a length of 200 bp and assigned to 32-nt bins along the genome. The resulting histograms (genomic “signal maps”) were stored in bigWig files. Narrow peaks were identified using the MACS 2.1.0 algorithm at a cutoff of p-value 10^-7^, without control file, and with the “–nomodel” option. Peaks that were on the ENCODE blacklist of known false ChIP-Seq peaks were removed. Signal maps and peak locations were used as input data to Active Motifs proprietary analysis program, which creates Excel tables containing detailed information on sample comparison, peak metrics, peak locations and gene annotations. For differential analysis, reads were counted in all merged peak regions (using Subread), and the replicates for each condition were compared using DESeq2 ^43^. Analysis was conducted using the following software: bcl2fastq2 (v2.20) for processing of Illumina base-call data and demultiplexing, Samtools (v0.1.19) for processing of BAM files, BEDtools (v2.25.0) for processing of BED files, wigToBigWig (v4) for generation of bigWIG files and Subread (v1.5.2) for counting of reads in BAM files for DESeq2 and HOMER motif analysis tool. ATAC-regions with |shrunken log_2_ FC| > 0.25 (IAV day 14 vs. PBS control) and an adjusted p-value < 0.1 were considered significant. Additional analysis and data visualization was performed using the Deeptools library ^44^, self-written Python scripts and IGV Genome Browser ^45^. Gene ontology analysis of gene loci with differential ATAC-regions was performed using the ClueGO plugin for Cytoscape Software. ATAC-seq data were deposited online and are available at NCBI’s Gene Expression Omnibus (GEO) under reference ID: GSE225498.

## Results

### Susceptibility to pneumococcal secondary infection depends on the serotype

In order to analyze sustained effects of influenza virus infection on AECII antibacterial responses, we employed a previously established murine infection model. Intranasal infection of C57BL/6 mice with a sublethal dose of a strictly pneumotropic, mouse-adapted strain of IAV (PR/8/34, H1N1) resulted in acute viral pneumonia ^17^. IAV-mediated disease included moderate, transient weight loss (Fig. 1A) and disruption of the pulmonary barrier leading to increased diffusion of blood-derived components (*i.e.* serum albumin) into the bronchoalveolar space (Fig. 1B). Of note, intraalveolar leakage of serum albumin was most pronounced during acute infection (day 7 post IAV) and significantly decreased during the post-IAV recovery phase. Yet, the pulmonary barrier was not fully restored at day 14 post infection (p.i.), as indicated by significantly elevated serum albumin levels in bronchoalevolar lavage fluid (BALF) from post-IAV mice (day 14) compared to control (PBS-treated) mice. At day 14 after IAV infection, post-IAV and control mice were infected with one of 3 strains of *Streptococcus pneumoniae* (*S.pn.*) of serotypes 4 (strain TIGR4), 7F (strain BHN54) and 19F (strain BHN100), all differing in their invasive disease potential (4 > 7F > 19F). Taking into account previously reported influenza-mediated changes in the upper respiratory tract of humans ^17, 46^ and mice ^47^, which might affect bacterial translocation into the lower airways, pneumococcal inocula were generally administered by oropharyngeal aspiration. At 18 h post pneumococcal infection, the airway bacterial burden was significantly increased during the post-IAV recovery phase for the *S.pn.* serotype 4 (Fig. 1C) and 7F (Fig. 1D) strains as compared to the respective *S.pn.* infection alone. In stark contrast, decreased airway CFU were detected already at 4 h as well as 18 h post *S.pn.* 19F infection in post-IAV mice (Fig. 1E). In line with increased airway bacterial loads of the *S.pn.* serotype 4 and 7F strains, a significantly increased incidence of bacteremia was observed after secondary *S.pn.* infection with these strains (Fig. 1F, 1G), as compared to bacterial infection alone, while *S.pn.* 19F generally did not cause bacteremia (Fig. 1H).

**Fig. 1:**
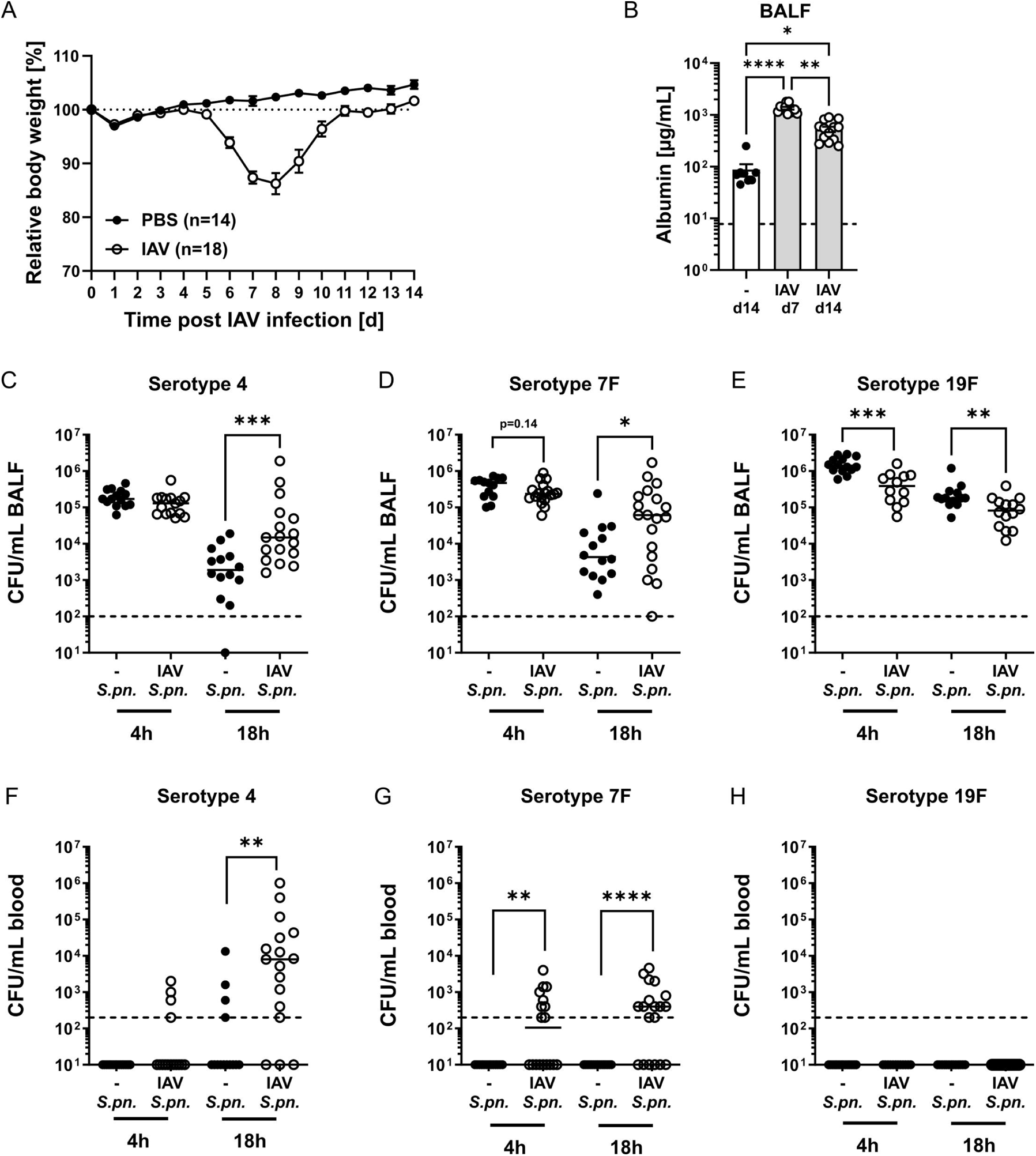
Influenza A virus (IAV) infection decreases lung barrier integrity and alters susceptibility towards *Streptococcus pneumoniae* (*S.pn.*). Mice were intranasally infected with 7.9 TCID_50_ IAV (H1N1, PR/8/34) or treated with PBS. **A)** Body weight of PBS- and IAV-treated mice. Mean % body weight (relative to baseline at d 0) ± SEM is graphed. Data were compiled from 3 independent experiments. **B)** Serum albumin in bronchoalveolar lavage fluid (BALF). Data were compiled from 2 independent experiments with n = 4-7 mice/group/experiment. Mean (bars) ± SEM and individual values are depicted. The dashed line indicates the detection limit. Statistical analysis was performed by Kruskal-Wallis and Dunn’s multiple comparisons test, *p>0.05, **p<0.01, ****p< 0.0001. At day 14 post primary treatment, IAV- and PBS-treated mice were oropharyngeally infected with 10^6^ *S.pn.* (serotype 4, 7F or 19F). Pneumococcal CFU in BALF **(C, D, E)** and blood **(F, G, H)** were assessed at 4 h or 18 h post bacterial infection. Statistical analysis was performed by two-tailed Mann-Whitney (C,D,E) or Fisher‘s exact test (F,G,H), *p<0.05, **p<0.01, ***p<0.001, ****p<0.0001.

Taken together, *S.pn.* bacteremic dissemination and outgrowth in the respiratory tract were enhanced in post-IAV mice, strongly depending on the *S.pn.* serotype. Serotype 4 showed the most invasive potential already in *S.pn.* single infection, with its invasivness drastically increasing in IAV-experienced hosts. Invasiveness of serotype 19F was not affected by preceeding IAV infection and even revealed lower CFU in this scenario. Serotype 7F, which in *S.pn.* infection alone was not invasive within 18 h of infection, managed however to cross the post-IAV lung mucosal barrier. This rendered especially the IAV/7F infection model interesting for exploring mechanistic clues to the synergism between primary IAV and secondary *S.pn.* infections with respect to IAV-mediated changes in AECII.

### The airway inflammatory microenvironment during secondary *S.pn.* infection is altered in a bacterial strain-dependent fashion

We next sought to determine whether the strain-dependent impaired clearance of *S.pn.* during post-IAV recovery (14 days post infection) would be associated with IAV-related and/or *S.pn*.-induced alterations in the airway inflammatory microenvironment. To this end, we performed bead-based immunoassay-profiling of 19 cytokines and chemokines in BALF from PBS-, IAV-, *S.pn.-* and IAV/*S.pn.*-treated mice at 4 h and 18 h post *S.pn.* infection. Analyte concentrations were z-score-transformed, hierarchically clustered and color-coded (Fig. 2A). Data from only the 4 h time-point are additionally presented in Fig. 2B to better visualize immediate-early responses.

**Fig. 2:**
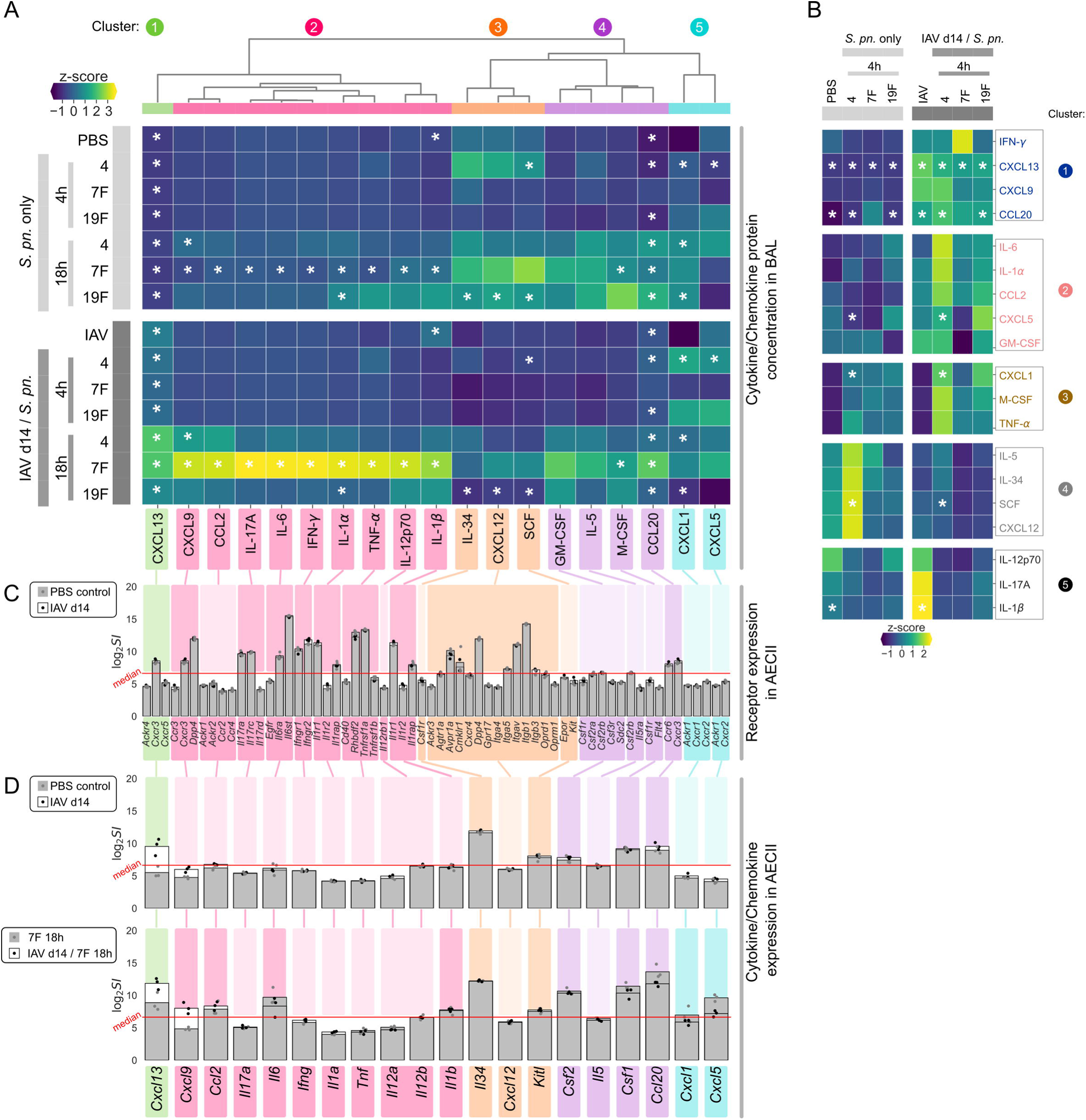
Resolved influenza A virus (IAV) infection alters airway cytokine and chemokine levels upon *Streptococcus pneumoniae (S.pn.)* infection. Mice were intranasally infected with 7.9 TCID_50_ IAV (H1N1, PR/8/34) or treated with PBS. On day 14 post infection (p.i.), IAV-infected and PBS-treated mice were oropharyngeally infected with 10^6^ *S.pn.* (serotype 4, 7F or 19F). Cytokine/chemokine concentrations in bronchoalveolar lavage fluid (BALF) were assessed at 4 h or 18 h post bacterial infection. Data were compiled from 1 - 3 independent experiments with n = 2 - 7 mice/group/experiment, such that each analyte was covered by 3 - 19 (average: 11) data points per group. **A)** Z-scores of mean cytokine/chemokine concentrations of each experimental group were used for heatmapping and clustering (cluster 1 - 5). Data for only the early (4 h) BALF cytokine/chemokine levels **(B)** are additionally shown. IAV/*S.pn.* conditions were compared to the respective *S.pn.* only condtion and IAV only condtion was compared to PBS control, with a two-sided Mann-Whitney-U test. Asterisks indicate * p < 0.05. **C)** Gene symbols of known receptors to clustered BALF cytokine/chemokines were obtained from the receptor/ligand database CellTalk DB. AECII receptor expression as log_2_ signal intensity (log_2_ SI) for PBS control and IAV 14 days p.i. was derived from AECII microarray analysis (n = 3 per group). Bars represent mean log_2_ SI with grey and black points indicating values of microarray replicates. Receptors with expression above median signal intensity of the microarray’s total SI-distribution (red line) are highlighted by the according BALF cytokine/chemokine cluster color. **D)** AECII expression levels of BALF cytokine/chemokine genes matching the respective receptors and cytokine/chemokine proteins in the BALF were derived from microarray data for 7F 18 h single-infection and IAV/7F 18 h secondary infection. Gene expression above median SI is highlighted by the according BALF cytokine/chemokine cluster color.

Comparison of BALF from PBS-*vs.* IAV only-treated mice revealed a small, but significant, increase in the inflammatory mediators CXCL13, IL-1β and CCL20 (Fig. 2D and supplementary figure S1).

Following *S.pn.* infection alone, highest cytokine/chemokine levels were observed after 18 h. While similar levels of GM-CSF, IL-5 and CXCL5 were detected in *S.pn.*-vs. IAV /*S.pn.-* infected airways, previous IAV-infection partly blunted induction of IL-34, CXCL12 and SCF upon secondary *S.pn.* infection with most serotypes (Fig. 2A). Immediate-early pro-inflammatory responses 4 h post IAV /*S.pn.-* infection showed subtle and partly strain-specific signatures (Fig. 2B, clusters 1-3). While resolved IAV infection was associated with significantly increased early (4 h) induction of CXCL13, CCL20, CXCL5 and CXCL1 upon secondary infection with *S.pn.* serotype 4 (Fig. 2B), it particularly favored the development of a hyperinflammatory microenvironment during secondary infection with *S.pn.* 7F at 18 h post pneumococcal infection (Fig 2A). This included significantly increased levels of several cytokines associated with type 1 (TNF-α: 12-fold increase, IL-12: 4-fold increase, IFN-γ: 123-fold increase) and type 3 (IL-17A: 20-fold increase) immune responses. Of note, IFN-γ showed the by far largest induction (Fig 2A). Also CXCL13, CXCL9, CCL2, IL-6, IL-1α, IL-1β, M-CSF and CCL20 showed significantly increased protein BALF concentrations after IAV/7F secondary infection (Fig 2A). These observations underlined the strain-specific impairment of bacterial clearance and containment following resolved IAV infection, that had been particularly evident for strain 7F (Fig 1).

By means of transcrptomic data, we next addressed the responsiveness of post-IAV AECII to the previuosly characterized airway cytokine/chemokine milieu. To this end, we isolated AECII from PBS-treated and post-IAV lungs (day 14) by flow-cytometry and analyzed transcription of the same 19 cytokines/chemokines and their corresponding receptor genes by microarray analysis and making use of the CellTalk receptor/ligand database.

Interestingly, gene expression of known receptors to measured BALF-cytokines/chemokines were similarly expressed in AECII cells, comparing resolved IAV infection with the PBS control. Yet, not each receptor gene analyzed was expressed above the median signal intensity level (average signal distribution is shown in supplementary figure S18 B). In this regard, AECII isolated following IAV infection and from control (PBS-treated) mice showed strong expression of receptors for CXCL13 (*Cxcr3*), CXCL9 (*Cxcr3* and *Dpp4*), IL-17a (*Il17ra* and *IL17rc*), IL-6 (*Il6ra* and *Il6st*), IFN-γ (*Ifngr1* and *Ifngr2*), IL-1α and IL-1β (*Il1r1* and *Il1rap*), TNF-α (*Rhbdf2* and *Tnfrsf1a*), CXCL12 (*Avpr1a*, *Cmklr1*, *Dpp4*, *Itga5*, *Itgav*, *Itgb1* and *Itgb3*) and CCL20 (*Ccr6*, *Cxcr3*), as shown in (Fig. 2C).

Next to cytokine/chemokine receptor expression, AECII are known sources for many of the BALF cytokines and chemokines previuosly analyzed on the protein level. We furthermore tested whether induction of cytokine/chemokine gene expression in AECII in response to *S.pn.* strain 7F (18 h post infection) was altered by previous IAV infection, as this *S.pn.* serotype had shown the strongest IAV-dependent BALF cytokine/chemokine response. At steady-state (day 14, PBS control, Fig. 2C, upper panel), AECII only expressed few of the tested BALF-mediators above median expression level (*Il34*, *Kitl*, *Csf2*, *Csf1* and *Ccl20*). Further, *Cxcl13* was the only gene that showed clearly increased expression in AECII following resolved IAV infection as compared to the PBS control (Fig. 2C, upper panel). Next to *Cxcl13*, only *Cxcl9* expression was clearly increased upon IAV /7F 18 h secondary infection as compared to 7F single infection. With respect to the hyperinflammatory BALF microenvironment detected 18 h following IAV/7F infection (Fig. 2A, lower panel), AECII therefore presumably made only minor contributions to soluble mediator production (Fig. 2C, lower panel) and protein mediators detected at high concentrations in the hyperinflammatory BALF cytokine/chemokine cluster 2 were likely of different cellular origin than AECII. Nevertheless, AECII do express receptors mediating responses to cluster 2 inflammatory mediators in secondary *S.pn.* infection. Further, we detected distinct changes in AECII gene transcription comparing resolved IAV infection with PBS treatment and *S.pn.* 7F infection with and without previous IAV infection (Fig. 2C). Therefore, we performed comprehensive analyses of the AECII transcriptional response towards all 3 *S.pn.* serotypes alone or in secondary infection following resolved IAV infection to develop mechanistic concepts for AECII responses to secondary *S.pn.* infections.

### ARACNE gene co-expression network analysis dissects AECII transcriptional response to interferons

Motivated by the unexpectedly distinct serotype-specific inflammatory responses detected in the BALF cytokine/chemokine profile especially following secondary *S.pn.* infection and the suprisingly stable expression of cytokine/chemokine receptors in AECII, we further analyzed their contribution to shaping serotype-specific cytokine responses in secondary *S.pn.* infection. We isolated vital lung AECII 4 and 18 hours post oropharyngeal *S.pn.* infection with serotypes 19F, 7F or 4 of previously IAV-infected (day 14) or PBS-treated (day 14) C57BL/6 mice. This resulted in triplicate AECII samples from 14 experimental conditions: **1.)** PBS, **2.)** IAV day 14, **3.)** serotype 19F 4 h, **4.)** serotype 19F 18 h, **5.)** serotype 7F 4 h, **6.)** serotype 7F 18 h, **7.)** serotype 4 4 h, **8.)** serotype 4 18 h, **9.)** IAV day 14 & serotype 19F 4 h **10.)** IAV day 14 & serotype 19F 18 h, **11.)** IAV day 14 & serotype 7F 4 h, **12.)** IAV day 14 & serotype 7F 18 h, **13.)** IAV day 14 & serotype 4 4 h, **14.)** IAV day 14 & serotype 4 18 h. Of note, *S.pn.* infected mice were treated with intranasal application of PBS 14 days prior *S.pn.* infection to reflect effects of the vehicle solution and application prodcedure. Gene transcription of AECII was analysed by microarray technology and differential gene expression was calculated in reference to AECII from PBS-treated mice. Between the 14 experimental conditions in total, 1,037 genes with an absolute transcriptional regulation fold-change of above 3 (versus PBS control AECII) and an ANOVA-based p-value < 0.05 were identified (supplementary table T1). To gain mechanistic clues to serotype-specific infection-induced AECII transcriptional responses, we constructed a gene co-expression network based on the “Algorithm for the Reconstruction of Accurate Cellular Networks” (ARACNE) from these genes ^31, 34, 48^. ARACNE aims at identifying direct transcriptional interactions by infering pair-wise mutual information (MI) between regulated genes and subsequent efficient pruning of distracting and likely indirect correlations, commonly arising in cascades of regulatory transcriptional interactions and their feed-back responses. Thereby, ARACNE generates a graph representation of the gene-regulatory network topology in which genes are represented as nodes that are connected by edges, representing potential direct regulatory interactions. Supplementary table T2 provides node and edge information of the resulting network.

ARACNE gene co-expression inference resulted in a network with 449 genes and 791 MI-based correlations between them (Fig. 3A). The five genes with the highest node connectivity k were transcription factors *Irf7* (k = 29, Interferon regulatory factor 7) and *Stat1* (k = 23, Signal transducer and activator of transcription 1) as well as the genes *B2m* (k = 18, Beta-2 microglobulin), *Epb41* (k = 16, Erythrocyte membrane protein band 4.1) and *Ly6a* (k = 13, Lymphocyte antigen 6 complex locus A). The network was partitioned into nine distinct modules (termed: M1 - M9, supplementary figure S2), with modules 1 and 2 containing most genes (M1: 139 genes, M2: 95 genes) and thus dominating the network. Dominance of M1 and M2 not only related to the number of genes but also to their intra- and inter-modular connectivity, as *e.g.* evidenced by the number of genes in M1 and M2 with intermodular connections (Fig. 3B). Of note, though M1 contained 44 genes more than M2, both modules had similar edge counts (M1: 217 edges, M2: 200 edges), indicating that genes in M2 were much more densely connected within their module than genes in M1.

**Fig. 3:**
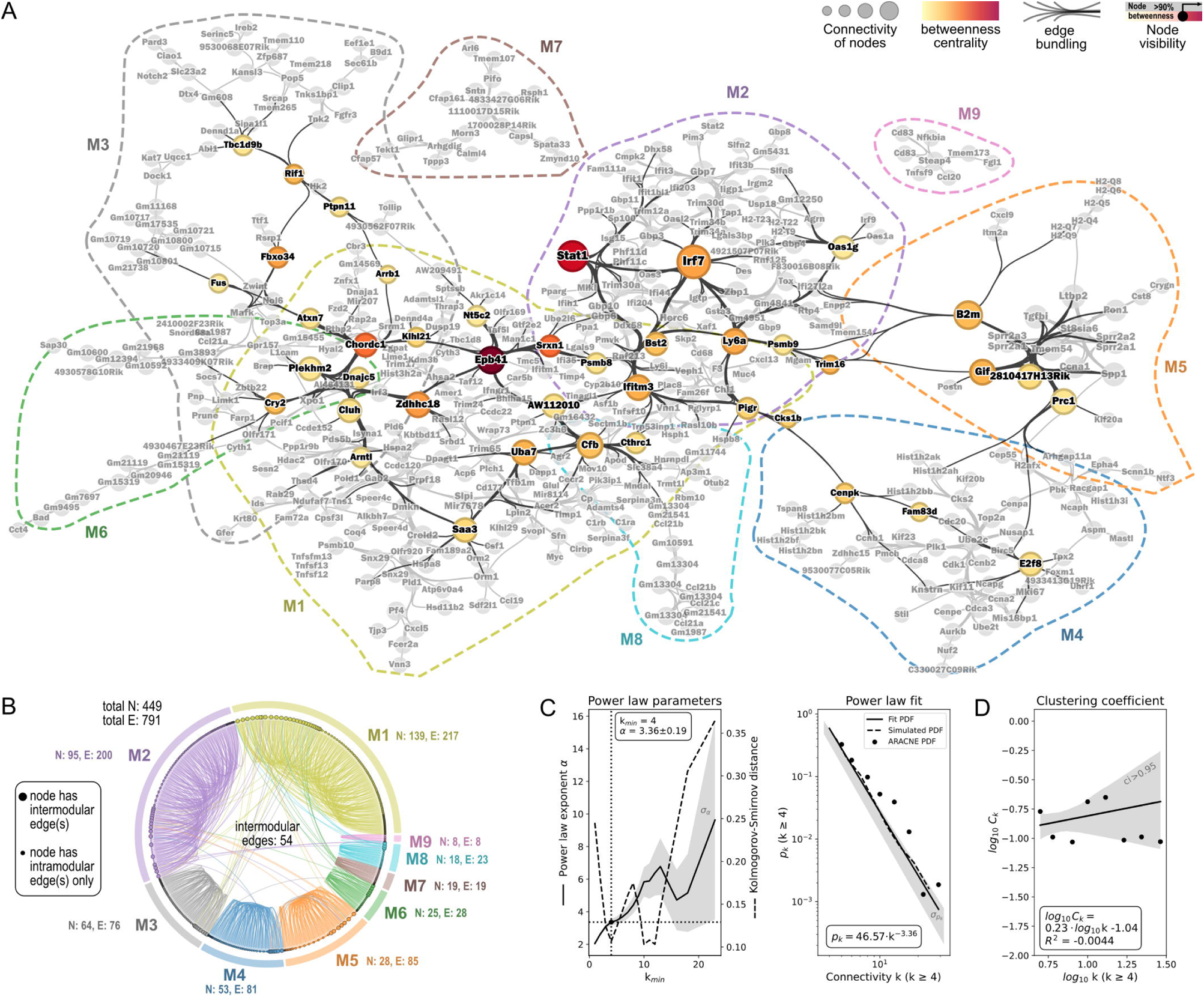
AECII ARACNE gene co-expression network inference. Mice were intranasally infected with 7.9 TCID_50_ IAV (H1N1, PR/8/34) or treated with PBS. On day 14, IAV-infected and PBS-treated mice were oropharyngeally infected with 10^6^ *S.pn.* (serotype 4, 7F or 19F). AECII were isolated 4 h or 18 h post bacterial infection from pooled lung cells of n = 3 -5 mice per group from 3 replicate experiments. The transcriptome was analysed by microarray analysis for differential gene expression between the mean expression per condition and the mean of the PBS control AECII. Genes with fold changes (FC) with |FC| > 3 in at least one condition were tested for significance by ANOVA (p<0.05). Normalized log_2_ SI data of all significantly differentially regulated genes were further analysed by the ARACNE gene co-expression network inference algorithm (see text for details). **A)** The resulting ARACNE network was partitioned using modularity/spectral partitioning into nine modules M1 – M9 (module outlines are indicated by colored dashed lines) and visualised followig multi-gravity force-atlas node positioning. The node size indicates node connectivity. Node labels indicate gene symbols. Edge visualization was simplified by the hammer edge bundling algorithm. Betweenness centrality of genes was caculated and genes with betweenness centrality values above the 90^th^ percentile are highlighted. Their betweenness centrality is color coded and their edges are shown in black. Remaining genes and edges are grayed out. **B)** Chord plot representation of the ARACNE network and modules. N: number of nodes, E: number of edges. Intra- and inter-modular edges are shown. **C)** Power law regression fit of node connectivity k. Left: Optimization of fit parameters k_min_ and power law exponent α based on Kolmogorov-Smirnov fitting quality. Right: Best power law fit of p_k_ vs. k for k ≥ 4. p_k_: Binned fractions of nodes with connectivity k. Gray areas represent variance (σ) of α and p_k_, respectively. **D)** Clustering coefficient distribution characteristic of nodes with k ≥ 4. C_k_: Binned fractions of nodes with clustering coefficient C and connectivity k. Gray area represents the 95 % confidence interval.

Figure 3A additionally gives a qualitative impression on the major hub genes of the network. This is based on the highest node betweenness centrality measure (> 90^th^ percentile). Betweenness centrality of a node is a measure of how many shortest paths between any pair of nodes in the network traverse that given node. Betweenness centrality was highest for *Epb41, Stat1, Srxn1* (Sulfiredoxin 1)*, Chordc1* (Cysteine and histidine rich domain containing 1)*, Zdhhc18* (Zinc finger DHHC-type palmitoyltransferase 18)*, Fbxo34* (F-box protein 34)*, Irf7, Ly6a, Ifitm3* (Interferon induced transmembrane protein 3) and *Trim16* (Tripartite motif containing 16), revealing these genes to play a central role in the AECII gene co-regulatory network.

We classified the obtained ARACNE network to be of scale-free-like nature, based on log_10_ p_k_ vs. log_10_ k power law fit of the node connectivity distribution (Fig. 3C, left). The optimal power law fit for k ≥ 4 (Fig. 3C, right) had an absolute value of 3.36 ± 0.19 for the exponent α, with genuine scale-free networks based on the Barabási–Albert model having power law exponents between 2 and 3 ^49, 50^. In line with this, the ARACNE network’s clustering coefficient distribution in the log_10_ C_k_ vs. log_10_ k plot was independent (R^2^ = -0.004) from the node connectivity (Fig. 3D), as can be expected for a network of this type ^51^. A key feature of scale-free network topology is that it relies on few heavily interconnected node hubs and many weakly interconnected peripheral nodes, rendering network responses resistent to randomly induced node failure or removal, but highly sensitive to hub node failures. Considering the scale-free-like topology of our ARACNE AECII-response network, in subsequent analyses we focused on the two apparent hub genes *Irf7* and *Stat1* with the highest node connectivities.

To understand how genes within the network’s nine modules were regulated in the different infection scenarios, their log_2_ fold change regulation, as compared to PBS control AECII were seperately clustered and color coded (Fig. 4A).

**Fig. 4:**
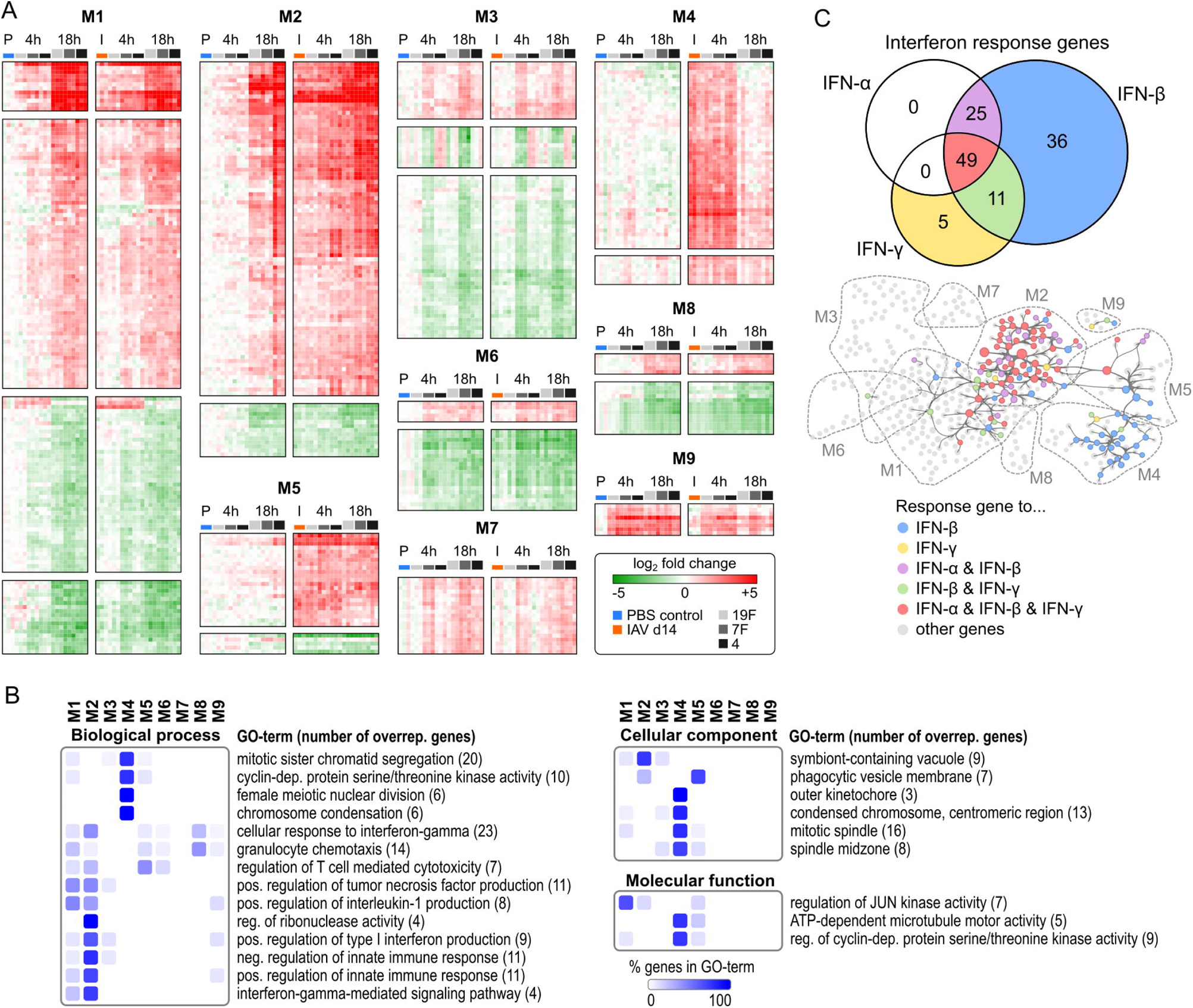
The ARACNE gene co-expression network reveals dominance of IFN-triggered responses in AECII in resolved IAV and acute secondary *S.pn.* infection. **A)** Clustered and color-coded log_2_ fold changes (over PBS control) of genes in the ARACNE network modules M1 – M9. Left sub-clusters represent PBS (day 14) treatment and *S.pn.* infections alone, right sub-clusters represent IAV infection (day 14) and secondary IAV/*S.pn.* infections with *S.pn.* serotypes 19F, 7F and 4. **B)** Gene ontology enrichment analysis (two-sided hypergeometric test, FDR < 0.05) of genes in the ARACNE network for indicated GO categories. Genes associated with significant GO-terms were mapped to ARACNE network modules (M1 – M9) and percent of term-associated genes per module were calculated, clustered and color-coded. The total number of network genes per GO-term is stated in brackets. **C)** Genes in the ARACNE network were analyzed for interferon (IFN-α, IFN-β, IFN-γ) response genes using the Interferome database. The numbers of genes responsive to a specific or multiple IFNs are presented as a Venn diagram. The groups of IFN-response genes, as displayed in the Venn diagram, were mapped to the ARACNE network. Dashed lines represent network module outlines.

Functional gene ontology (GO) annotation of genes in the nine modules (Fig. 4B) clearly attributed M1 and even more M2 to interferon responses and innate immune responses as well as IL-1 and TNF production. Interestingly, M1 contains several gene products regulating Jun kinase activity, a biochemical mechanism induced by stress responses leading to assembly of the JUN/FOS (AP-1) transcription factor complex ^52, 53^. Previously, we observed unique responses to secondary infection with serotype 7F with respect to bacterial clearance and inflammation. Interestingly, M3 emerged in the ARACNE network as an especially 7F-related module. However, this module was hardly associated with common GO-terms, so that its functional implications remain unclear. M4 clearly attributed to mitosis, proliferation and chromatin organisation. Cellular component GO-terms showed genes in M5 to be related to phagocytic vesicles, while M8 was to some extent pointing at granulocyte chemotaxis. Like M3, also the small modules M6, M7 and M9 did not clearly annotate to any common GO-terms, likely due to their low gene count. Nevertheless, a number of genes from M9 (e.g. *Ccl20, CD83* and *Nfkbia*) encode for well known players in the context of acute inflammation ^54–56^.

As GO-term enrichment suggested a major involvement of the interferon system in the AECII response, we additionally mapped all ARACNE network nodes to type I (IFN-α, IFN-β) and type II (IFN-γ) interferon response genes (Fig. 4C) using the Interferome database^57^. We set a 3-fold regulation cutoff and limited the response time-frame to 18 h to as much as possible to match the Interferome database search with our dataset. We found that out of the 449 genes in the ARACNE network, 126 (28 %) are listed in the Interferome database as responders to type I and/or type II interferon signaling events. It is known that the different interferons induce partly overlapping transcriptional responses, hence 49 out of the 126 IFN-response genes (∼ 39 %) respond to IFN-α, IFN-β as well as IFN-γ. Of note, 36 genes (∼ 29 %) specifically respond to IFN-β only. Only 5 genes were exclusive IFN-γ response genes (∼ 4 %) and none of the genes were specific to IFN-α. Out of the 126 IFN-response genes, 25 (∼ 20 %) were shared between IFN-α and IFN-β and 11 genes (∼ 9 %) between IFN-β and IFN-γ (Fig. 4C top). Mapping the individual contributions of interferon-response genes to the previously infered ARACNE network revealed a dominant impact of interferon response genes on the co-expression network. This was clearly centered within M2, but also reached into M1, M4, M5 and M9 (Fig. 4C, bottom). Of note, M2 IFN-response genes were largely related to all three considered interferons. However, IFN-response genes in M4 seemed to be inducible only by IFN-β and otherwise annotated to cell proliferation, based on GO-enrichment. This indicated that IFN-β might be relevant for AECII proliferation or replenishment following pathogenic encounter.

Taken together, we successfully inferred an ARACNE-based scale-free-like gene co-expression network from transcriptional profiles of AECII sorted from PBS-treated, only *S.pn.*-infected (three serotypes), only IAV-infected and IAV/*S.pn.*-infected (three serotypes) mice, encompassing immediate-early (4 h p.i.) and acute (18 h p.i.) transcriptional responses, respectively. This network revealed distinct hub-genes in the AECII transcriptional response and was dominated by two modules (M1 and M2), with M2 appearing heavily related to interferon response genes.

### Serotype-specific transcriptional AECII-responses are related to differential co-expression network usage

To i) comprehend how the topological ARACNE gene co-expression network structure and its modularity relate to the temporal regulation of gene transcription in AECII upon *S.pn.* infection alone and secondary *S.pn.* infection following IAV infection and ii) to identify serotype-dependent differences in network usage, we mapped FC regulation (in reference to PBS control AECII) to the ARACNE network for all experimental conditions. To this end, network nodes were color coded according to the log_2_ fold change regulation of the respective transcript at a given experimental condition. Additionally, network nodes and their adjacent edges were hidden if the represented gene did not show significant regulation with |FC| > 3. For reference, Fig. 5A shows the assignment of nodes to the nine network modules M1 – M9. Figure 5B shows the network representations for AECII isolated from only *S.pn.* infected mice (19F, 7F, 4; 4 h and 18 h p.i.). Figure 6 represents the resulting network maps for AECII isolated following IAV infection alone (day 14, Fig 6A) and secondary *S.pn.* infection following IAV infection (IAV/19F, IAV/7F, IAV/4; 4 h and 18 h p.i.; Fig 6B). Supplementary figures S5 - S15 provide high-resolution network representations with full gene symbol labeling.

**Fig. 5:**
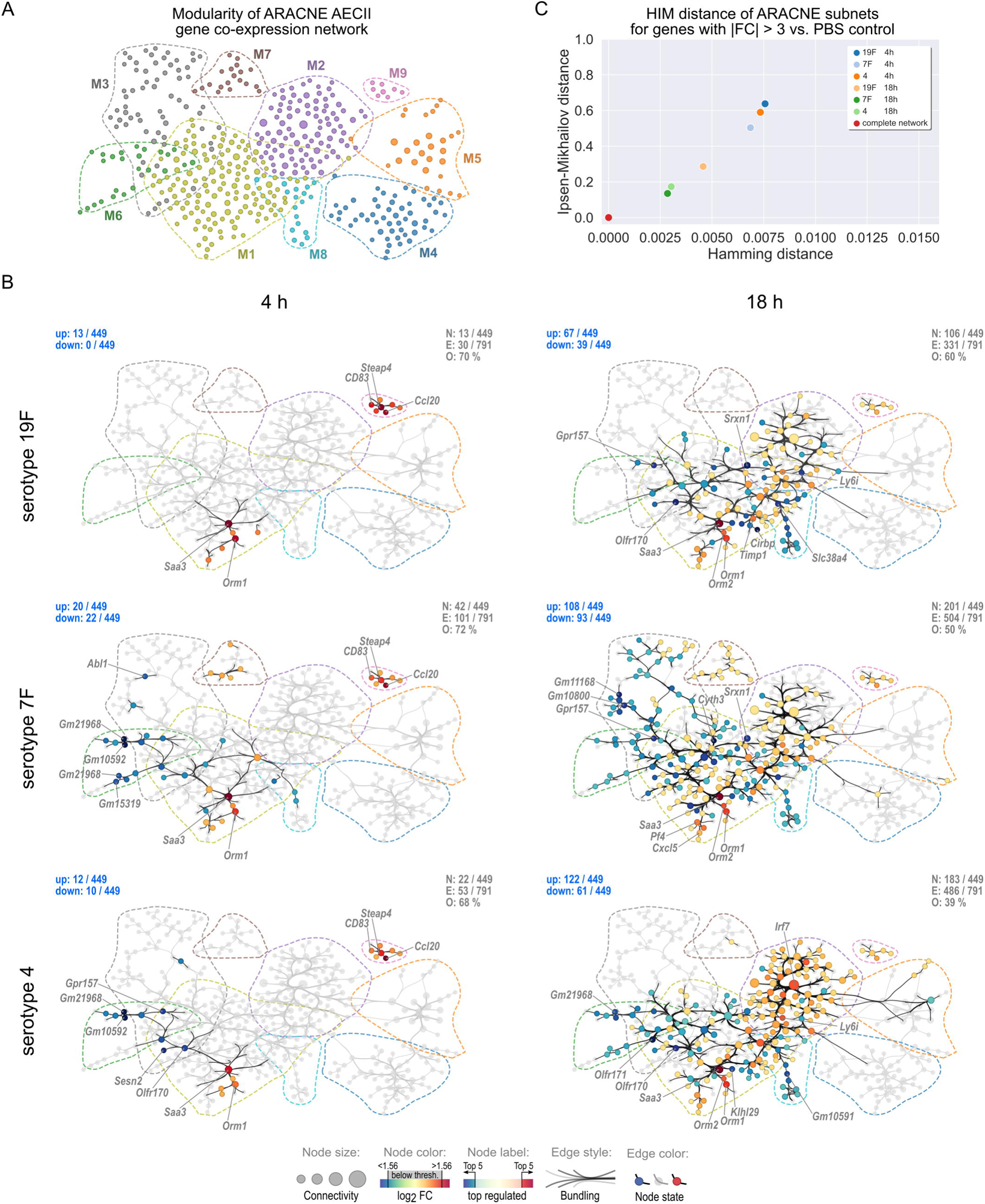
AECII ARACNE differential gene co-expression partial networks reveal serotype-specific network response patterns following *S.pn.* infection alone. **A)** Module assignment of individual genes to ARACNE network modules M1 – M9. Dashed lines indicate module outlines. **B)** Partial networks for genes with |FC| > 3 and their associated edges were calculated for *S.pn*. infections with the indicated serotypes alone at 4 h and 18 h post infection. Nodes are color-coded according to log_2_ FC. Remaining nodes and edges are grayed-out. Node size indicates node connectivity. Dashed lines indicate outlines of network modules. Gene symbols of the top 5 most intensely up/down regulated genes are labeled. N: number of nodes with |FC| > 3 vs. PBS control. E: number of highlighted edges. O: Percent of egdes that are greyed-out. Numbers of up/down-regulated genes are stated in blue in the upper left corners. **C)** HIM distances of FC-based partial networks shown in B in reference to the complete ARACNE network.

**Fig. 6:**
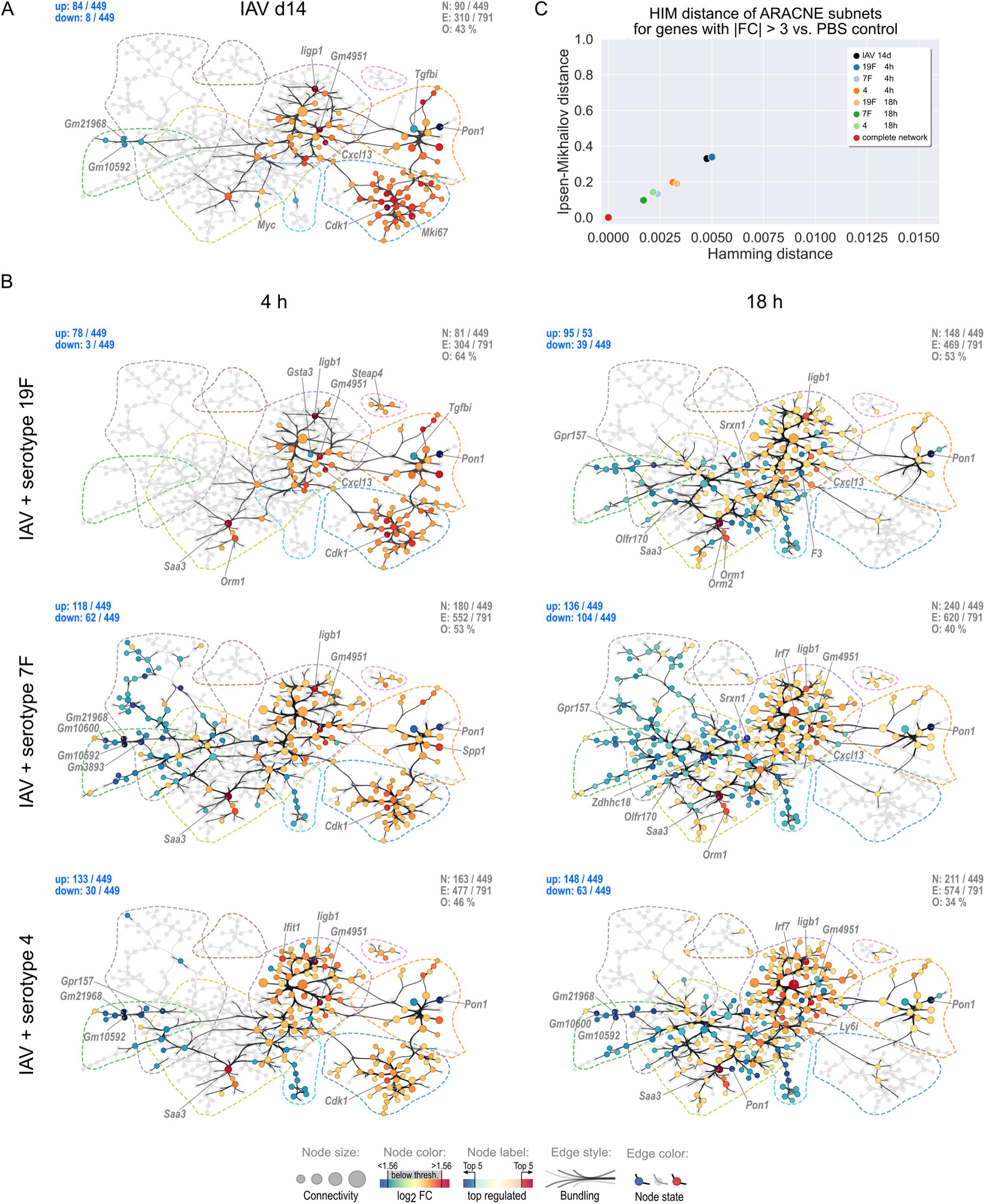
AECII ARACNE differential gene co-expression partial networks reveal IAV-mediated changes in AECII transcriptional regulation and serotype-specific network patterns in secondary *S.pn.* infection. Partial networks for genes with |FC| > 3 and their associated edges were calculated for the AECII gene transcriptional profile detected 14 days post IAV infection **(A)** and 4 h and 18 h following secondary *S.pn.* infection with serotypes 19F, 7F or 4 **(B)**. Nodes are color-coded according to log_2_ FC. Remaining nodes and edges are grayed-out. Node size indicates node connectivity. Dashed lines indicate outlines of network modules. Gene symbols of the top 5 most intensely up/down regulated genes are labeled. N: number of nodes with |FC| > 3 vs. PBS control. E: number of highlighted edges. O: Percent of egdes that are greyed-out. Numbers of up/down-regulated genes are stated in blue in the upper left corners. **C)** HIM distances of FC-based partial networks shown in B in reference to the complete ARACNE network.

To compare topological features of the partial networks in Fig. 5B and Fig. 6B, defined by statistically significant at least 3-fold gene regulation for the respective experimental condition, we calculated the Hamming-Ipsen-Mikhailov (HIM) distance of each partial network in reference to the complete ARACNE network (Fig. 5C and 6C). The Hamming distance (H) of two networks evaluates the presence/absence of matching network edges, based on the difference of the two network adjacency matrices and thus reflects local network similarities ^58^. In contrast, the Ipsen-Mikhailov (IM) distance is a measure of global topological similarities between two networks, based on spectral comparisons of their adjacency matrices ^59^.

Interestingly, we found that as early as 4 h after *S.pn.* infection alone, serotype-specific patterns amongst the AECII differential partial networks started to emerge (Fig. 5B, left). Genes in M9 (*Ccl20*, *Steap4*, *CD83*, *Nfkbia, Tmem173*, *Fgl1* and *Tnfsf9*) became swiftly upregulated following infection with each serotype, with their expression reducing at 18 h p.i. Also, in M1 *Saa3* and *Orm1* were strong early transcriptional responders 4 h after infection with any of the three tested *S.pn.* serotypes. However, in M6, genes such as *Gm10592*, *Gm21968* (both predicted genes) and *Gpr157* were strongly downregulated in AECII only following 7F and serotype 4 infection. Also, 7F induced upregulation of genes in M7 (e.g. *Arhgdig*, *Arl6*, *Glipr1* and *Calml4*) 4 h p.i., which was not the case for serotypes 19F and 4. At 4 h post infection however, AECII generally did not show intense differential gene regulation of genes with |FC| > 3 within the ARACNE network (19F: 13 up; 7F: 20 up, 22 down; 4: 12 up, 10 down). This was also reflected in the HIM distances of the partial networks. Here, the three serotype-specific partial networks (4 h post infection) located farthest from the complete network and cumulated relatively close to one another (Fig. 5C).

The AECII transcriptional response to *S.pn.* infection alone drastically increased 18 h p.i. as compared to 4 h p.i. (19F: 67 up, 39 down; 7F: 108 up, 93 down; 4: 122 up, 61 down) and further encompassed serotype-specific network patterns. Transcriptional activity in M1 greatly enhanced around genes previously active (4 h). This related to both up- and down-regulation. Interestingly, 7F infection downregulated genes in M3, which was a unique feature of this serotype. Also, most genes in M6 were still downregulated 18 h p.i., which was however not the case for infection with the 19F serotype. In contrast, most genes in M8 (e.g. *Ccl21a*, *Ccl21b*, *Ccl21c*) were consistently downregulated in infections with all serotypes. At 18 h p.i., a major contribution to the structure of the three serotype-specific partial networks however was made by M2, containing mostly interferon response genes. The invasive serotype 4 *S.pn.* strain clearly triggered the strongest response in this module, inducing strong upregulation of virtually each node in the module. As compared to serotype 4, both, the number of genes regulated above threshold as well as the extent of their regulation were not as prominent for infection with serotypes 19F and 7F. M2 is dominated by interferon response hub transcription factors *Irf7* and *Stat1* that were highly expressed in all three partial networks 18 h post infection with any of the three *S.pn.* serotypes.

Following *S.pn.* infection alone, modules M4 and M5 were not extensively perturbed. In contrast, transcriptional perturbation of genes within M4 and M5 was a strikingly distinct feature of the late inflammatory aftermath (day 14 p.i.) following IAV infection alone (Fig. 6A). Here, AECII upregulated all genes in M4, which were previously found to be related to IFN-β responses and various mitotic mechanisms (Fig. 4B and C). Amongst the most upregulated genes from M4 were *e.g. Cdk1*, *Mki67*, *E2f8*, *Cdc20* and *Ube2c*, most of which are directly related to an AECII proliferative transcriptional configuration. M5 contains MHC class I (*B2m*) and II genes and similarly showed strong up-regulation of most accordingly attributed genes (*e.g. H2-Q5*, *H2-Q6*, *H2-Q7*, *H2-Q8*, *H2-Q9*, *Spp1*, *Prc1* and *Gif*), with the exception of *Tgfbi* and *Pon1*, that both were strongly downregulated. Of note, genes in M5 feature the highest clustering coefficients (supplementary figure S16) amongst all genes in the network, indicative of intricate correlative connections amongst its constituents. Importantly, many interferon response genes in M2 were also upregulated (e.g. *Cxcl13*, *Iigp1*, *Gm4951*, *Usp18*, *Trim30a*, *Bst2*, *Ly6i*, *Ly6a* and *Irf7*) in AECII on day 14 following IAV infection. Of note, transcriptional upregulation of *Cxcl13* 14 days post IAV infection was well in line with elevated Cxcl13 BALF protein levels (Fig. 1D). In M1, only few genes were up-regulated above threshold (*Rasl10b*, *Saa3*, *AW112010* and *Timp1*) and the important transcription factor *Myc* (c-Myc) was downregulated. In M6, transcription of four predicted genes (*Gm10600*, *Gm21968*, *Gm10592* and *Gm3893*) previously observed to be downregulated in *S.pn.* infection with serotypes 7F and 4 alone were also downregulated 14 days post IAV infection alone. In total, 14 days post IAV infection, 84 out of 449 ARACNE network genes were up-regulated (> 3-fold) and 8 genes were down-regulated. The regulated genes in total had 310 (of 791) MI-based correlations, positioning the IAV (day 14) partial network at about the same HIM-distance as the partial networks of the *S.pn.*-only infections 18 h p.i. (Fig. 6B and 5B). Importantly, these results indicated resolved IAV infection to sustainably affect the transcriptional profile of AECII in the lung. This finding underlined our rationale to study AECII-responses towards *S.pn.* in secondary infection following resolved IAV infection.

The IAV-shaped transcriptional network configuration provided the basis for the AECII response to secondary *S.pn.* infection with serotypes 19F, 7F and 4 (Fig. 6B). At 4 h following secondary *S.pn.* infection with serotype 19F, the number of at least 3-fold regulated genes and the resulting partial network remained similar to the IAV-shaped partial network. In line with this, the HIM distances of the IAV (day 14) and the IAV/19F (4 h) conditions were similar (Fig. 6C). In contrast, infection with serotypes 7F and 4 resulted in a swift enlargement of the AECII partial co-expression network, positioning them much closer to the complete network and setting them clearly apart from the IAV/19F (4h) partial network in the HIM distance plot. In particular, secondary infection with serotypes 7F and 4 rendered the regulated interferon response genes in M2 more numerous as compared to IAV/19F infection (4 h). Of note, the comparative blunting of the immediate early AECII response to secondary infection with serotype 19F may be linked to the reduced BALF CFU observed in this infection

At 18 h post secondary pneumococcal infection, resolved IAV infection clearly enhanced the transcriptional regulation of interferon response genes in M2 for all serotypes as compared to the respective *S.pn.* infection alone. This effect was most pronounced for the 7F serotype (also compare Fig. 4A), which became particularly evident when focusing on the two major interferon-regulatory hub genes *Irf7* and *Stat1* and their network neighbors (supplementary figure S17). The uniqueness of the AECII response 18 h post IAV/7F secondary infection was reflected also by the HIM distance of the according partial network, which was closest to the complete ARACNE network as compared to all other conditions (Fig. 6C).

As for *S.pn.* infection alone, serotype 7F also exclusively induced downregulation of most genes attributed to M3 in secondary infection following IAV infection, only now already 4 h post *S.pn.* infection.

Interestingly, most proliferation-associated genes in M4 underwent a major gene regulatory shift 18 h post secondary *S.pn.* infection with any serotype, reversing the proliferative transcriptional configuration of AECII detected 14 days post IAV infection alone. Similarly, many genes in the adjacent module M5 became downregulated as well, again counteracting the IAV-primed induction of this module.

Taken together, ARACNE network inference of the *in vivo* AECII transcriptional response to different serotypes of *S.pn.* alone or following resolved IAV infection allowed elaborate dissection of serotype-specific responses. These analyses identified a positive transcriptional synergy between sustained IAV-mediated alterations to AECII gene transcription and *S.pn.* infection, leading to an accelerated and intensified interferon response, particularly in case of serotype 7F. This finding interestingly co-incided with the previously observed enhanced susceptibility of IAV-infected mice to invasive disease and excess inflammation following secondary infection with *S.pn.* 7F. In contrast, previous IAV infection had little effect on the transcriptional upregulation of interferon response genes to serotye 4 infection, which *per se* showed their strongest induction as compared to serotypes 7F and 19F. Next to enhanced, synergistic AECII responses with respect to IFN-signalling, secondary *S.pn.* infection led to an abrogation of IAV-associated proliferation and presumably repair mechanisms in AECII irrespective of the pneumococcal serotype.

### Resolved IAV-infection alters the airway IFN-response towards *S.pn.* and the responsiveness of AECII to IFNs

We sought to examine, whether the observed changes in the post-influenza AECII response to *S.pn.* and especially in their IFN-signalling would be associated with an altered abundance of airway IFN in secondary *S. pn.* infections. Thus, we compared levels of bioactive IFN I and III in BALF from PBS-treated as well as IAV-, *S.pn*.- and IAV/*S.pn.* infected mice using type I and III IFN-sensitive epithelial cells isolated from Mx2-Luc reporter mice ^30^. Airway IFN I/III levels in post-IAV (day 14), 7F as well as 19F-infected (18 h) mice were similar to baseline airway IFN levels in PBS-treated controls. Infection with *S.pn.* serotype 4 strain was associated with elevated airway IFN responses, independent of previous IAV infection. In contrast, previous IAV infection was associated with significantly increased IFN I/III-bioactivity upon infection with the *S.pn.* 7F strain as compared to 7F infection alone (Fig. 7A). Prompted by these findings, we sought to determine which specific IFNs were associated with the serotype-specifically altered inflammatory responses to pneumococci in post-influenza lungs. To this end, we quantified IFN-α, -β, -λ2/3 and -γ by ELISA and bead-based immunoassay, respectively. Similar to the results obtained by IFN-bioassay, previous IAV-infection greatly enhanced IFN-α responses in the airways upon *S.pn.* 7F infection, reaching similar IFN-α levels as compared to *S.pn.* serotype 4 infection (Fig.7B). Interestingly, we further observed that, as compared to baseline (PBS control, day 14), airway IFN-β was reduced 14 days following IAV infection alone and not boosted in infection with either *S.pn.* serotype 4, 7F nor 19F. In contrast, secondary infection with *S.pn.* 7F following IAV infection enhanced IFN-β responses as compared to 7F alone, yet only to concentrations that were comparable to baseline levels in the PBS-treated control group (Fig. 7C). In contrast to type I IFN production, induced by either *S.pn.* and/or previous IAV infection, we only detected minor changes in type III IFN. Here, only infection with *S.pn.* 7F alone was associated with a small, but significant decrease of IFN-λ2/3 (Fig. 7D). Lastly, and in partial analogy to the serotype-specific effects of previous IAV infection on antipneumococcal type I IFN, we observed a very strong and significant increase in airway IFN-γ when comparing *S.pn.* infection alone to secondary *S.pn.* infection exclusively for 7F (Fig. 7E).

**Fig. 7:**
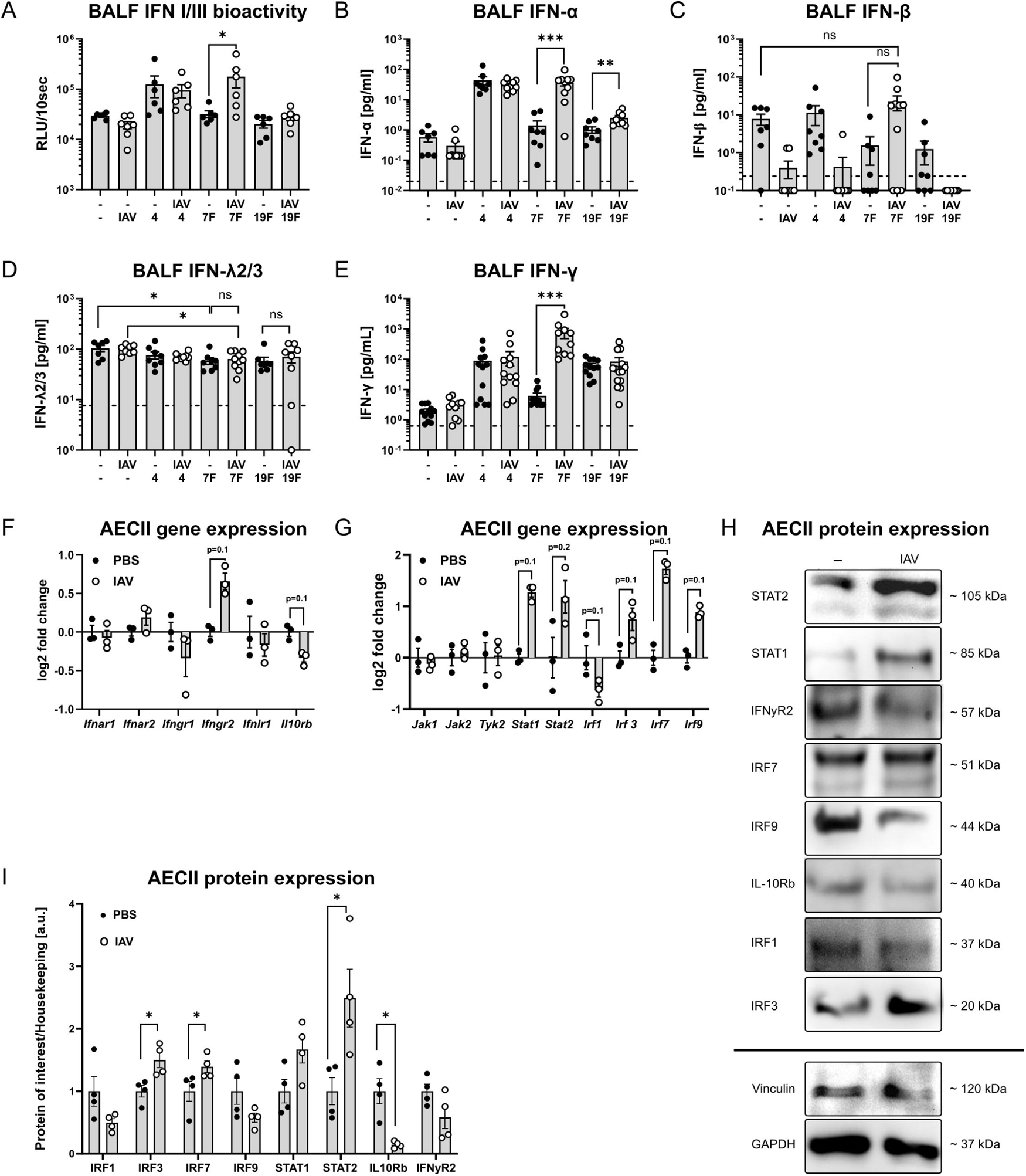
IAV-mediated alterations in airway IFN protein levels and AECII expression of IFN-signaling-related genes. **A-E)** Mice were intranasally infected with 7.9 TCID_50_ IAV (H1N1, PR/8/34) or treated with PBS. At day 14, IAV-infected and PBS-treated mice were oropharyngeally infected with 10^6^ *S.pn.* (serotype 4, 7F or 19F). Levels of bioactive type I/III IFNs **(A)**, IFN-α **(B)**, IFN-β **(C)**, IFN-λ **(D)** and IFN-γ **(E)** were determined in BALF 18 h post pneumococcal infection using an Mx2-luc-reporter assay **(A)**, ELISA **(B, C, D)** or bead-based multiplex assay **(E)**. Data were obtained from at least two independent experiments. Statistical analyses were performed by 2-way ANOVA and Tukey‘s multiple comparisons test, *p<0.05, ***p<0.01, ***p<0.001. **F-I)** Gene transcription and protein expression in AECII were analyzed by microarray or western blot. AECII from n = 3-5 mice/experimental group/experiment were pooled. Depicted are individual data and mean ± SEM from at least 3 independent infection experiments. Statistical analysis was performed by two-tailed Mann-Whitney test, *p<0.05. **F)** Relative expression of IFN receptor genes. **G)** Relative expression of IFN signal transducer genes. **H)** Representative blots from AECII protein lysates. **I)** Relative expression of selected IFN-signaling proteins, normalized to the expression of the housekeeping proteins vinculin or GAPDH.

Next to alterations in respiratory IFN-levels, it is conceivable that IAV-mediated intrinsic changes in the AECII responsiveness to IFNs contributed to their altered gene transcriptional response to IFNs in secondary *S.pn.* infection. Therefore, we analyzed the expression of molecules involved in IFN-detection and - signalling in AECII isolated from either PBS-treated control lungs or post-IAV lungs (day 14) in the previously generated microarray data and western blot analyses, respectively. We detected increased gene expression of *Ifngr2* at day 14 post IAV, whereas expression of *Il10rb* was decreased (Fig. 7F). Of note, dedicated interrogation of the microarray data for AEC interferon expression itself of any kind revealed expression (in descending strength) of *Ifna5*, *Ifnl3*, *Ifnl2*, *Ifna9*, *Ifna15*, *Ifna6* and *Ifna1* above median level (supplementary figure S18) wich was remarkably stable across all infection conditions.

Notably, expression of various intracellular interferon signal transducers (*Stat1*, *Stat2*, *Irf3*, *Irf7*, *Irf9*) was increased in AECII after IAV infection, while only expression of *Irf1* was reduced (Fig. 7G). Except for IFNGR2 and IRF9, we found that changes observed on mRNA levels could be similarly detected on the protein level (Fig. 7I, H and Supplementary figure S19).

Taken together, AECII isolated from IAV convalescent mice 14 days post infection generally expressed increased levels of IFN-response transcription factor proteins STAT2, IRF7 and IRF3 as compared to AECII from uninfected mice. This cellular state likely contributed to the characteristics of *S.pn.* serotype 7F secondary infection in convalescent IAV-infected mice and enabled an enhanced AECII response to type I and II interferons, as compared to serotypes 19F and 4. This is fully in line with core observations of the previous AECII transcriptome analysis.

### Epigenetic imprinting of AECII MHC-class II expression and IFN-responses

The sustained presence of an IAV-associated transcriptional footprint in AECII made it conceivable that this was at least partly mediated via epigenetic imprinting. *E.g.* epigenetic imprinting of AECII transcriptional responses could underly the faster and stronger response of AECII to interferons upon secondary *S.pn.* encounter. Therefore, we analyzed AECII from IAV-infected (day 14) and PBS control mice by ATAC-seq to identify genomic regions with differential chromatin accessibility (Fig. 8). In total, we identified 83,681 ATAC-regions (of which 78 % were annotatable) within AECII chromatin (from two independent replicates) over both conditions (supplementary table T3). From these, 260 regions showed significant differential ATAC-activity (|shrunken log_2_ FC| > 0.25, FDR < 0.1). 118 of the 260 regions showed enhanced and 142 regions decreased chromatin accessibility (Fig. 8A). Enhanced regions were systematically shorter (median: 670 bp) than depressed regions (median: 818 bp). *De novo* HOMER motif-enrichment analysis indicated significant over-representation of 3 sequence motifs (Fig. 8A bottom). The main motifs GTGACTCA (33 sites) and CACATTCCTA (16 sites), found within ATAC-regions with enhanced accessibility after IAV-infection, are known to be compatible with JUN/FOS (AP-1) and TEAD (TEAD1, 2 and 4) transcription factors, respectively. ATAC-regions with impaired accessibility related to the family of forkhead-binding-domain-containing transcription factors with the main motif GCAAACACTG (20 sites). Of note, *Foxm1* was one of the forkhead-motif-binding transcription factors also part of the AECII ARACNE network (M4), providing further clues to the consequences of diminished chromatin accessibility of according forkhead-motif sites. GO annotation (biological process) of gene loci positioned within a broad anotation window arround differential ATAC-regions showed association with type I interferon signaling and production (*Chuk, Zbtb20, Oas1e, Oas3, Nlrc5* and *Slamf8*), nucleosome assembly (*Hat1, H2bc7, H2bc6, H4c4, Smyd3* and *H2bc8*) and vesicle endocytosis (*Itsn1, Ap2b1, Syt2* and *Lrrk2*) (Fig. 8B).

**Fig. 8:**
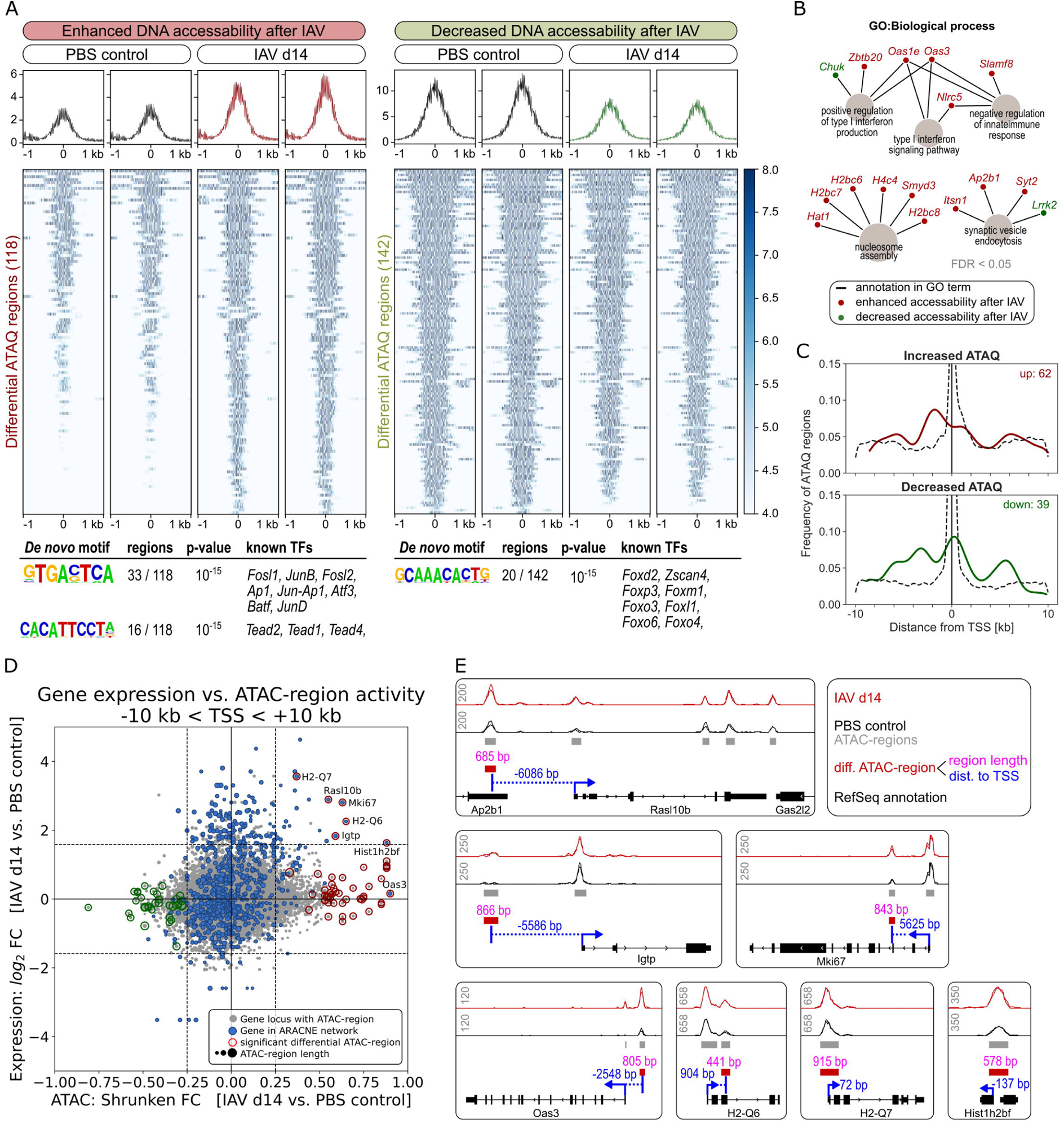
IAV-infection induces epigenetic imprinting of MHC-class II and interferon response genes in AECII. Mice were intranasally infected with 7.9 TCID_50_ IAV (H1N1, PR/8/34) or treated with PBS. After 14 days, AECII were isolated from n = 3 -5 pooled mice per experimental group in two replicate experiments and chromatin was analyzed by ATAC-seq. DNA regions with differential ATAC-state (|shrunken log_2_ FC| > 0.25, FDR < 0.1) comparing IAV day 14 AECII vs. PBS control AECII were identified. **A)** More (118) and less (142) active ATAC-regions were aligned at region centers and normalized ATAC-signals within a window of ± 1 kb were clustered and color-coded. Significant results of HOMER analysis for transcription factor binding motifs and known binding TFs are indicated below. **B)** Gene Ontology (GO) analysis (two-sided hypergeometric test, FDR < 0.05) of genes annotating to differentially active ATAC-regions for the GO category “biological process”. **C)** Histogram showing relative localization of differentially active ATAC-regions. Only regions with gene annotations that fall into a window of ± 10 kb around transcription start sites (TSS) were included. Dashed lines represent localization of all ATAC-regions with positive and negative shrunken FC without thresholding, respectively. **D)** Scatterplot correlating genes with at least one differentially active ATAC-region within a window of ± 10 kb of TSS and respective microarray gene expression data. Each grey point represents an annotated ATAC-region. Point size represents ATAC-region length. Microarray-based log_2_ FC of gene symbols are plotted vs. ATAC-based shrunken log_2_ FC of matching ATAC-regions. ATAC-regions annotating to gene symbols that are part of the ARACNE network are shown in blue. Significantly less/more active ATAC-regions are indicated by green and red circles, respectively. **E)** Locus representation of genes of the ARACNE network that yielded significant differential activity of ATAC-regions. Data represent overlaid ATAC-signals from two independent replicates of AECII from PBS controls and resolved IAV infection (day 14). Grey numbers in upper left corners indicate normalized bigwig-data scaling.

As expected, on the genome-wide scale, the 83,681 ATAC-sites in AECII were mainly positioned around transcription start sites (TSS), as shown in Fig. 8C (dashed lines). Constriction of the differential ATAC-region annotation window to ±10 kb around adjacent TSS limited the number of relevant sites to 62 sites with significantly enhanced ATAC-activity and 39 sites with reduced activity, respectively. Interestingly, ATAC-sites with enhanced accessibility were favoured to be positioned in proximal promoters up to -4 kb upstream of TSS, whereas most sites with decreased accessibility were positioned around TSS (Fig. 8C).

To match results from ATAC-seq analysis of AECII 14 days after IAV-infection with the ARACNE gene co-expression network, we next combined both approaches based on fold-differences over the PBS control. Fig. 8D shows a scatterplot that represents shrunken log_2_ ATAC-region fold changes of all identified ATAC-sites (gray points; significance of region accessibility is indicated by red/green circles) positioned in a window of ±10 kb around genetic start coordinates of known annotated murine gene loci alongside with the matching log_2_ fold change of transcript expression in the AECII microarray dataset (IAV day 14 vs. PBS control). This approach led to the identification of seven gene candidates (*Hist1h2bf*, *Igtp*, *Mki67*, *Rasl10b*, *H2-Q6*, *H2-Q7* and *Oas3*) that were part of the ARACNE network and at the same time yielded an ATAC-region with enhanced accessibility in the vicinity of their locus start. Transcript expression of all candidates with the exception of *Oas3* was significantly elevated in AECII 14 days post IAV infection over the PBS control. These findings suggested that in AECII, enhanced expression of *Hist1h2bf*, *Igtp*, *Mki67*, *Rasl10b*, *H2-Q6* and *H2-Q7* following IAV-infection relies on an epigenetic regulatory component.

Fig. 8E shows locus ATAC-states of all these gene candidates. *Igtp* (interferon gamma-induced GTPase) and *Oas3* (2’-5’-oligoadenylate synthetase 3) belong to the interferon response genes in ARACNE module M2 and were both positively correlated with the hub transcription factor *Irf7*. *Ras10lb* (RAS like familie 10 member b), a predicted G protein activator, is part of M1 and was one of few genes with intermodular connections within the ARACNE network, connected to *Ifitm3* in M2 and *B2m* in M5. *H2-Q6* and *H2-Q7* belong to M5 and were not connected to any adjacent transcription factors but were interconnected with other major histocompatibility genes in the same network module. *Mki67* (marker of proliferation Ki-67) in M4 is connected to transcription factors *E2f8* and *Foxm1*, both related to cell proliferation. *Hist1h2bf* (also termed *H2bc7* or H2B clustered histone 7) is also part of M4 and correlated to other histones from the H2b family.

Taken together, ATAC-analyses of AECII isolated 14 days post IAV infection identified six ARACNE network genes that were at the same time transcriptionally upregulated and revealed ATAC-regions with enhanced chromatin accessibility in their promoter regions. This finding provides valuable insights into epigenetic imprinting of AECII transcriptional responses after the resolution of IAV infection and their embedding into the AECII gene co-expression network.

## Discussion

We have previously described rapidly induced and exceptionally strong transcriptional regulation in AECII following *in vivo* IAV infection ^23^. At the same time, we have observed prolonged susceptibility towards *S.pn.* following IAV infection in a *S.pn.* serotype-specific manner ^17^. Prompted by these and other observations, we now hypothesized i) AECII to serotype-specifically respond to *S.pn.* infection *in vivo* and ii) IAV infection to sustainably alter *S.pn*.-directed AECII-responses, contributing to enhanced susceptibility towards secondary *S.pn.* infection. In a comprehensive experimental approach, we isolated murine primary AECII following IAV infection alone and at different time-points post *S.pn.* infection alone and secondary *S.pn.* infection following IAV infection. Analysis of AECII gene transcription was then performed to infer an AECII-centric ARACNE gene co-expression network. Comparing three serotypes of *S.pn.* (19F, 7F, 4) harboring different levels of invasive disease potential allowed us to address the *S.pn.* serotype-specific inflammatory lung milieu and bacterial clearance alongside with AECII responses represented by the ARACNE network’s node, connectivity structure and its differential usage. With respect to the AECII response to different *S.pn.* serotypes alone, we detected robust regulation of gene transcription that indeed revealed serotype-specific characteristics. This included up- and downregulation with varying kinetics and revealed serotype-specific differences already 4 h post infection. While IFN I and III have primarily been associated with viral infections, IFNs play a role also in anti-bacterial defense. In the AECII transcriptional response to *S.pn.* infection alone, the interferon hub transcription factors Stat1 and Irf7 were highly induced in response to all three serotypes. Serotype 4 further stood out with an exceptionally strong induction of the genes within the respective ARACNE module (M2). This was well in line with highest BALF IFN-α and -β levels in response to serotype 4 infection alone as compared to 19F and 7F alone (Fig. 7B, C). As IFN genes themselves were not particularly regulated in AECII (supplementary figure S18), this finding underlines the inflammatory milieu established by the plethora of cells present in the infected airways to shape AECII responses in respiratory infections.

Resolved IAV infection led to higher lung bacterial burdens following secondary infection with serotypes 4 and 7F and increased invasiveness of these serotypes as compared to *S.pn.* infection alone. This contrasted significantly improved bacterial clearance of serotype 19F from the airways in secondary infection as compared to 19F infection alone. Since IAV convalescence particularly favored bacterial outgrowth and systemic dissemination of *S.pn.* serotypes 7F and 4 but not 19F, increased invasiveness of *S.pn.* following IAV infection does not seem to be a general host-dependent phenomenon and is thus at least partially serotype-specific. Moreover, resolved IAV infection, within a timeframe of 18 h post secondary infection, lowered the threshold for invasive serotype 7F infection as compared to 7F infection alone. In line with this, immediate-early lung cytokine/chemokine responses were uniquely boosted only to 7F secondary infection (18 h) and were clearly different from those observed upon serotype 4 and 19F secondary infection.

In this context, IFN-γ showed the largest induction upon secondary 7F infection, a fact that is likely of relevance for the phenomenon of decreased immunity to *S.pn.* 7F following IAV infection. IFN-γ has previously been shown to impede host defense by suppressing the phagocytic capacity of alveolar macrophages (AMs) during post-IAV recovery ^10^. Of note, while enhanced susceptibility to secondary bacterial infections, as we observed it here for serotypes 7F and 4, is a widely accepted phenomenon, there is also a report of protection from pneumococcal serotype 4 infection 28 days following post IAV infection ^60^. Therefore, next to *S.pn.* serotype-related factors, the timing of secondary pneumococcal infection during recovery from IAV infection may play an equally critical role in determining the ultimate outcome of secondary *S.pn.* encounters.

With respect to the bacterial factors shaping serotype-specific inflammation, AECII responses and bacterial clearance, the pneumococcal capsule represents a strong candidate. It is the key determinant of individual pneumococcal serotypes, a prerequisite for invasiveness and thereby a major virulence factor ^61^. Only encapsulated strains can establish disease, while at the same time, production of capsule components involves energetically costly mechanisms ^12^. In this context, it has been assumed that bacterial nutrient availability in the airways is affected by IAV infection and contributes to enhanced susceptibility to secondary bacterial infection following IAV infection ^62^ and recently, capillary leakage during IAV infection has indeed been shown to provide nutrients and antioxidants supporting *S.pn.* outgrowth ^7^.

Despite serotype-specific manifestations, the AECII response to secondary infection with a particular serotype qualitatively and quantitatively never equaled the response to infection with that serotype alone. This demonstrates general IAV-associated modulation of anti-pneumococcal AECII responses independent of the *S.pn.* serotype (or in fact, possibly even the bacterial pathogen) encountered. Generally, there was a strongly intensified transcriptional regulation of IFN response genes in AECII following secondary *S.pn.* infection, with an especially intensified response in the case of secondary 7F infection (compared to 7F infection alone). IAV-mediated alterations in the secondary response to *S.pn.* were detected on top of sustained changes detected in AECII 14 days post IAV infection alone, a time point when viral clearance was completed. Importantly, IAV-mediated changes in AECII were evident both on the gene transcriptional as well as the epigenetic level, supporting the concept of inflammatory imprinting.

Acute inflammatory sensing of interferons by the respective interferon receptors ultimately induces a transcriptional response involving a dedicated set of interferon-stimulated genes (ISG). Canonically, phosphorylated transcription factor complexes composed of STAT1/STAT2 homo- and heterodimers in combination with further transcription factors like IRF1 and IRF9 shape the quality and quantity of ISG transcriptional responses. Given that 14 days post IAV infection, BALF levels of IFN-α, IFN-β and IFN-γ were comparable to or even lower than in the PBS control group (Fig. 7B,C,E) it may surprise that AECII anyhow continue to express a robust set of interferon response genes (e.g. in M2, Fig. 6A), accompanied with significantly elevated protein levels of IRF1 and STAT2 (Fig. 7H, I) in AECII. Although we did not assess the phosphorylation status of STAT-proteins, this nevertheless may hint *e.g.* to an altered IFN-detection threshold due to sensitized IFN-signaling cascades, or to a long-lasting transcriptional ISG locus memory that, to some extent, is independent of classical ISG-induction mechanisms *e.g.* via phosho-STAT protein complexes. As this seems to lay a molecular mechanistic foundation for the accelerated and intensified transcriptional AECII response to interferons upon secondary *S.pn.* infection, persistence of IAV-primed ISG expression in AECII is worth considering. Indeed, there is evidence for IFN-independent and -dependent roles of non-phosphorylated STAT1 and STAT2 for basal and long-term ISG expression ^63^, thereby providing a concept for ISG expression in a non-canonical fashion. This takes place without the necessity for a cellular environment constantly providing high levels of interferons, especially during the tissue recovery phase. In this regard, it is also important to note that the described IAV-mediated alterations were detected at a time-point when viral clearance was completed, thereby excluding IAV remnants as a trigger for late IFN-induction ^17^.

Using an imiquimod model of skin inflammation and the analysis of epidermal stem cells, deep mechanistic insight into inflammatory memory on the cellular level, that might also apply to explain persistent ISG expression in post-IAV AECII, has recently been achieved ^64^. In their model, Larsen *et al.* demonstrate that cell-type and stimulus-specific transcription factors in conjunction with common stress-response transcription factors FOS-JUN (also known as AP-1) manage to set genomic memory domains. Their associated gene loci take part in a later transcriptional inflammatory memory recall. According to this concept, JUN eventually remains bound to the initially generated genomic memory domains and together with further homeostatic transcription factors is responsible for an accessible chromatin state of memory domains. In case of a secondary inflammatory stimulus, stress-induced FOS is re-recruited independently of the initially required stimulus-specifc transcription factors and ultimately induces fast and efficient transcription of genes associated to the memory domains ^64^. As ATAC analysis of AECII 14 days post IAV infection identified 33 genomic regions with enhanced chromatin accessibility that contain binding motifs for JUN/AP-1 transcription factors, the model suggested by Larsen *et al.* may also apply to AECII. This provides a further mechanistic explanation for the persistent ISG expression that is likely to accelerate and enhance the AECII ISG response in secondary *S.pn.* infection. In line with this, ARACNE network module M1 contains 5 genes (*Ccl19*, *Dnaja1*, *Dusp19*, *Fzd2* and *Ptpn1*) that are GO-annotated to regulate JUN kinase activity, with JUN phosphorylation being biochemically important for JUN activation. However, further studies are required to *de facto* link chromatin-accessibility of ATAC-identified AP-1 sites in IAV-imprinted AECII to the expression of the observed set of persistently expressed IFN-response genes.

Another *S.pn.* serotype-independent phenomenon observed upon secondary infection with all three serotypes was the swift blunting of genes in the AECII proliferation-related network module M4. This module’s transcriptional activity is clearly a consequence of post-IAV tissue repair processes. Interestingly, IFN-α, IFN-β and particularly IFN-λ were demonstrated to impair AECII proliferation in regenerating mouse lungs post IAV infection, which has also implications for the severity of *S.pn.* secondary infection ^18^. Consequently, IFNLR-knockout mice show improved post-IAV AECII proliferation and are less susceptibile to *S.pn.* secondary infection ^18^. Given our finding of persistent ISG expression in AECII post IAV infection, the swift shutdown of the AECII mitotic transcriptional configuration may be a direct consequence of an accordingly increased AECII responsiveness to interferons during secondary *S.pn.* infection.

Since AECII act as progenitor cells to constantly replenish AECI pneumocytes ^19^ that thereby inherit epigenetic footprints from their cellular AECII ancestors, cumulatively our findings may also help to understand long-term inflammatory imprinting of respiratory mucosa pneumocytes, other than AECII.

In this study, we confirm that IAV infection sustainably predisposes mice to secondary *S.pn.* infection in a serotype-specific manner. Importantly, we show that in this context AECII sustainably retain an IAV/inflammation-triggered transcriptional configuration with epigenetic involvement. This is accompanied by faster and more vigorous AECII responses to interferons in secondary *S.pn.* encounter and has implications for AECII proliferation. Thereby we show that AECII need to be considered as potent contributors to inflammatory tissue imprinting in the respiratory tract following IAV infection and other triggers.

## Acknowledgements

The authors wish to thank colleagues at Helmholtz Centre for Infection Research: Lothar Gröbe and Maria Höxter for cell sorting as well as Petra Hagendorff, Martina Grashoff, Tatjana Hirsch, Regina Lesch, Hanna Shkarlet, Silvia Prettin and Karin Lammert for excellent technical assistance in microarray processing, interferon assays and mouse experiments, respectively.

## Author Contributions

JDB designed and performed animal experiments, AECII cell isolation, sample and assay processing and wrote the manuscript. AJ analyzed transcriptomics and ARACNE network, interpreted ATAC-Seq data and wrote the manuscript. KSZ performed animal experiments. LM performed western blots. KST performed animal experiments. EG provided help in ATAC data visualization, AK provided cell lines and protocols for interferon response assays. SSK and DB were responsible for conceptualization of the study, scientific recommendations and revised the manuscript.

## Supplementary Information

**Supplementary table T1:**

Complete microarray expression dataset.

**Supplementary table T2:**

Node and edge list of ARACNE network.

**Supplementary table T3:**

Identified ATAC regions.

**Supplementary Figure S1:**

Airway cytokine and chemokine levels in S.pn. and IAV/S.pn. infection.

**Supplementary Figure S2:**

Module assignment of ARACNE network genes.

**Supplementary Figure S3 – S15:**

Differential AECII ARACNE gene co-expression partial networks.

**Supplementary Figure S16:**

Clustering coefficients of ARACNE network nodes.

**Supplementary Figure S17:**

Clustered node neighbour expression heatmaps of interferon-hub-genes *Irf7* and *Stat1*.

**Supplementary Figure S18:**

Expression of interferon genes in AECII.

**Supplementary Figure S19:**

Western blot quantification of IFN-signaling-related proteins in AECII.

