## Supplementary Figures S1-S18 for "Epigenetic changes and serotype-specific interferon-responses of lung epithelial cells in late post-influenza pneumococcal pneumonia"

**Supplementary Figure S1: Airway cytokine and chemokine levels in *S.pn.* and IAV/*S.pn.* infection.** Mice were intranasally infected with 7.9 $TCID_{50}$ IAV (H1N1, PR/8/34 strain) or PBS. 14 days post primary treatment IAV- and PBS-treated mice were oropharyngeally infected with $10^6$ *S.pn.* (serotype 4, 7F or 19F). Cytokine/chemokine concentrations in bronchoalveolar lavage fluid (BALF) were assessed at 4 h or 18 h post bacterial infection by multiplex bead-based immunoassay. Data were compiled from 1 - 3 independent experiments with n = 2 - 7 mice/group/experiment. Z-scores of mean cytokine/chemokine concentrations of each experimental group were calculated. IAV/*S.pn.* conditions were compared with the according *S.pn.* only conditions and IAV only condtion was compared to PBS control with a two-sided Mann-Whitney-U test. * $p < 0.05$.

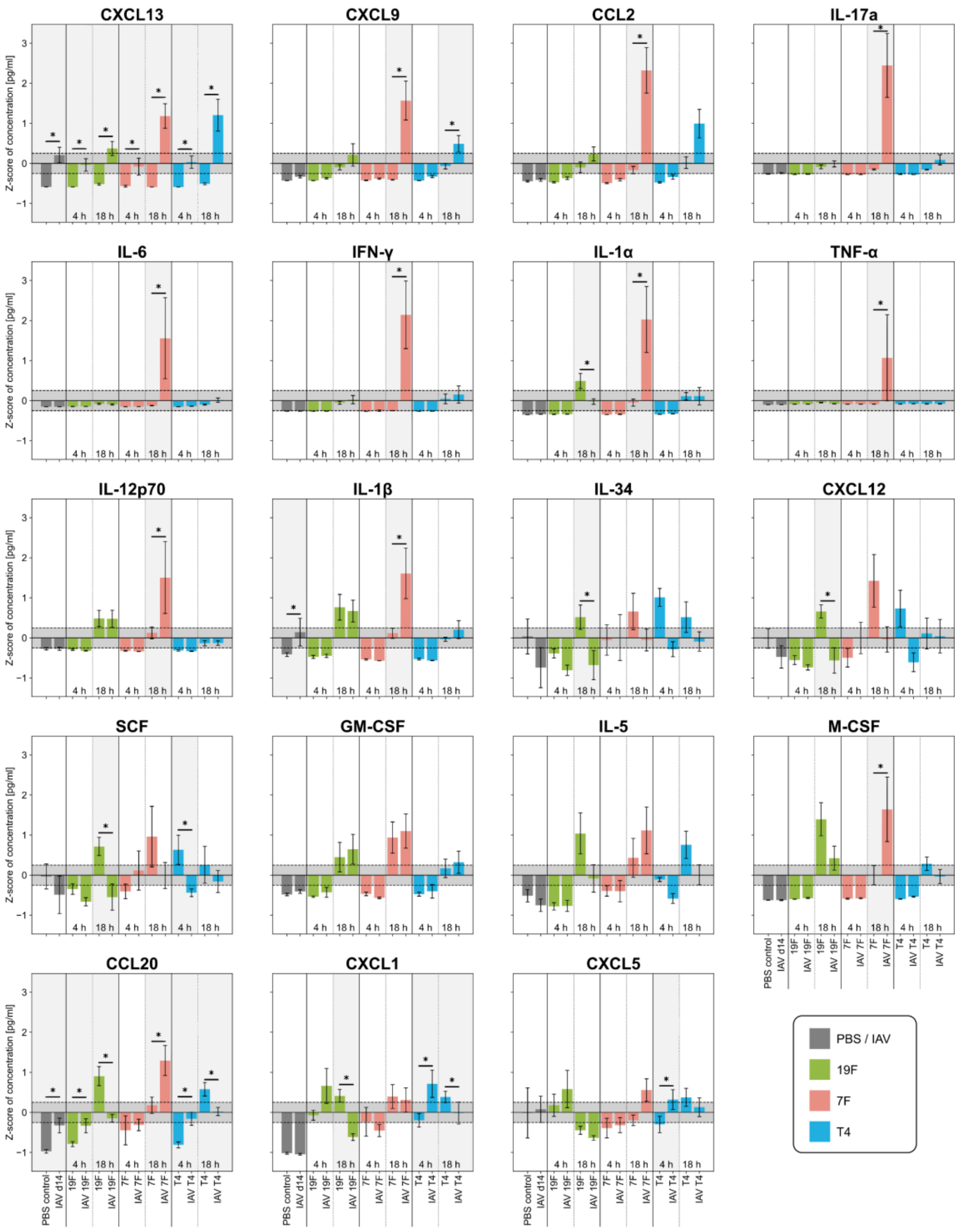

**Supplementary Figure S2: Module assignment of ARACNE network genes.** Module assignment of individual genes to ARACNE network modules M1 - M9. Dashed lines indicate module outlines. Nodes are labeled with gene symbols.

#### S2: Module assignment of ARACNE network genes

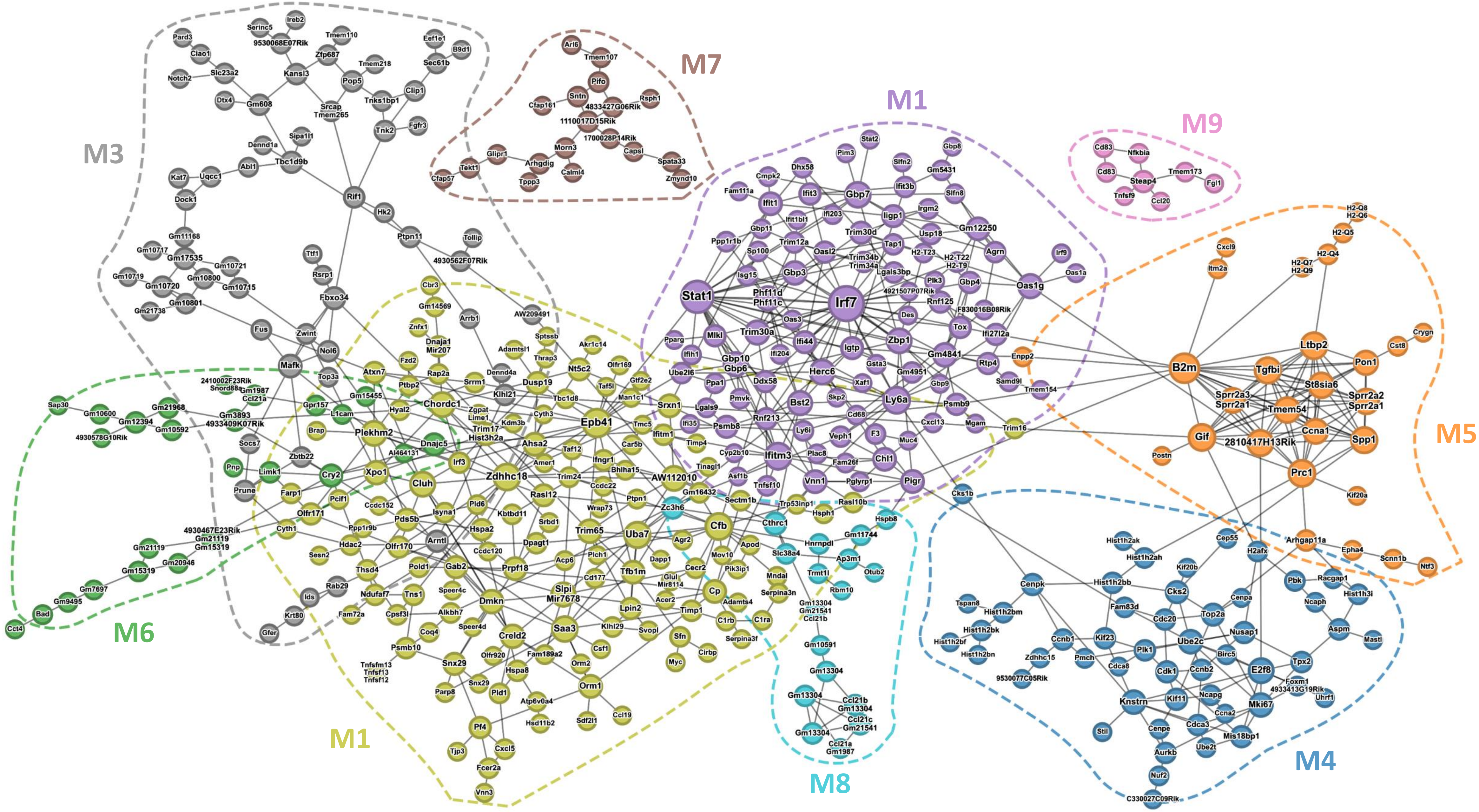

**Supplementary Figure S3 - S15: Differential AECII ARACNE gene co-expression partial networks.** Partial networks for genes with |FC| > 3 and their associated edges were calculated for all 13 infection conditions, stated in the upper right corner. Nodes are color-coded according to $\log_2$ FC. Remaining nodes and edges are greyed-out. Node size indicates node connectivity. Colored dashed lines indicate outlines of network modules. Nodes are labeled with gene symbols.

**S3: Serotype 19F 4 h vs. PBS control**

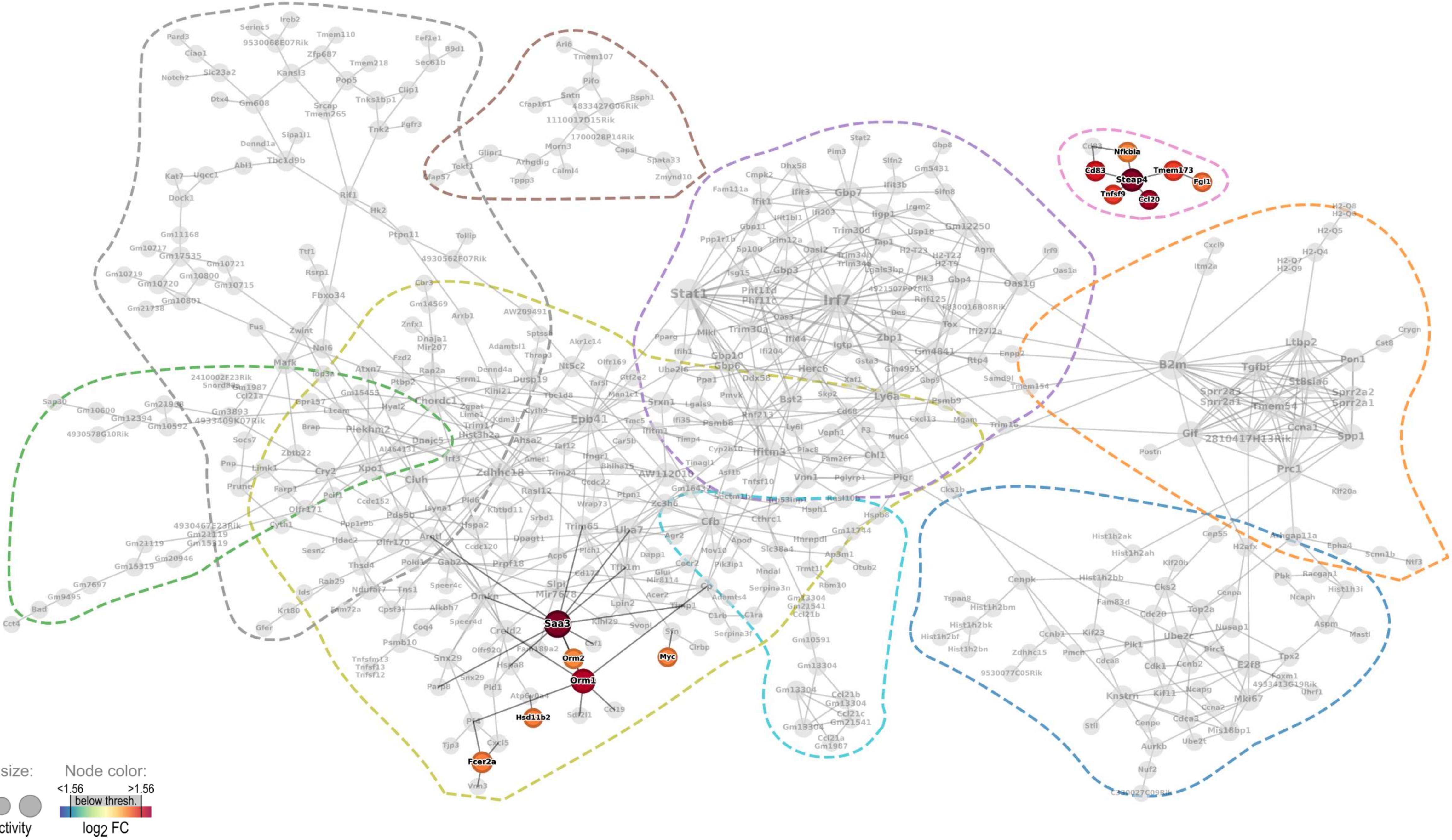

**S4: Serotype 7F 4 h vs. PBS control**

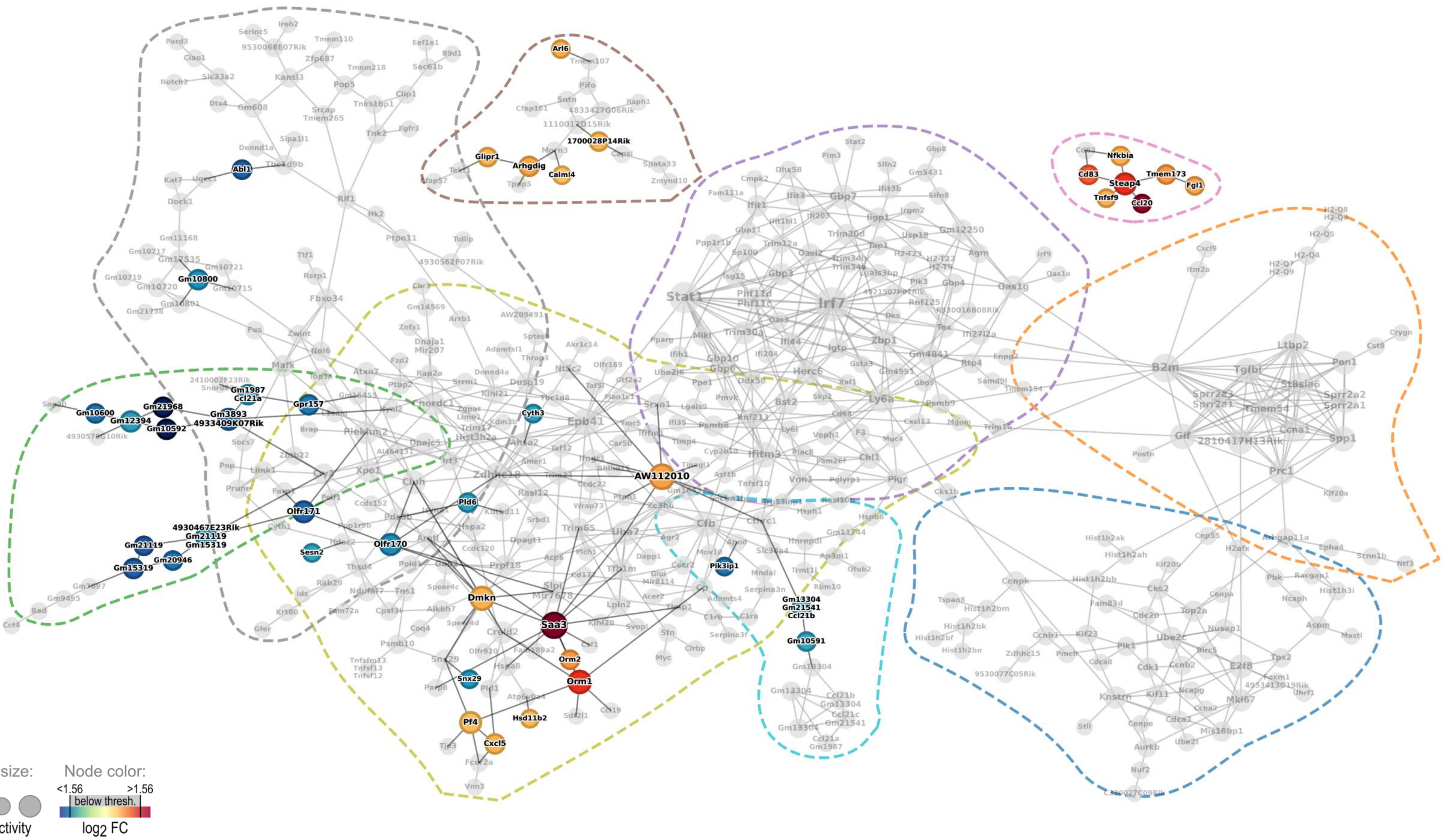

#### S5: Serotype 4 4 h vs. PBS control

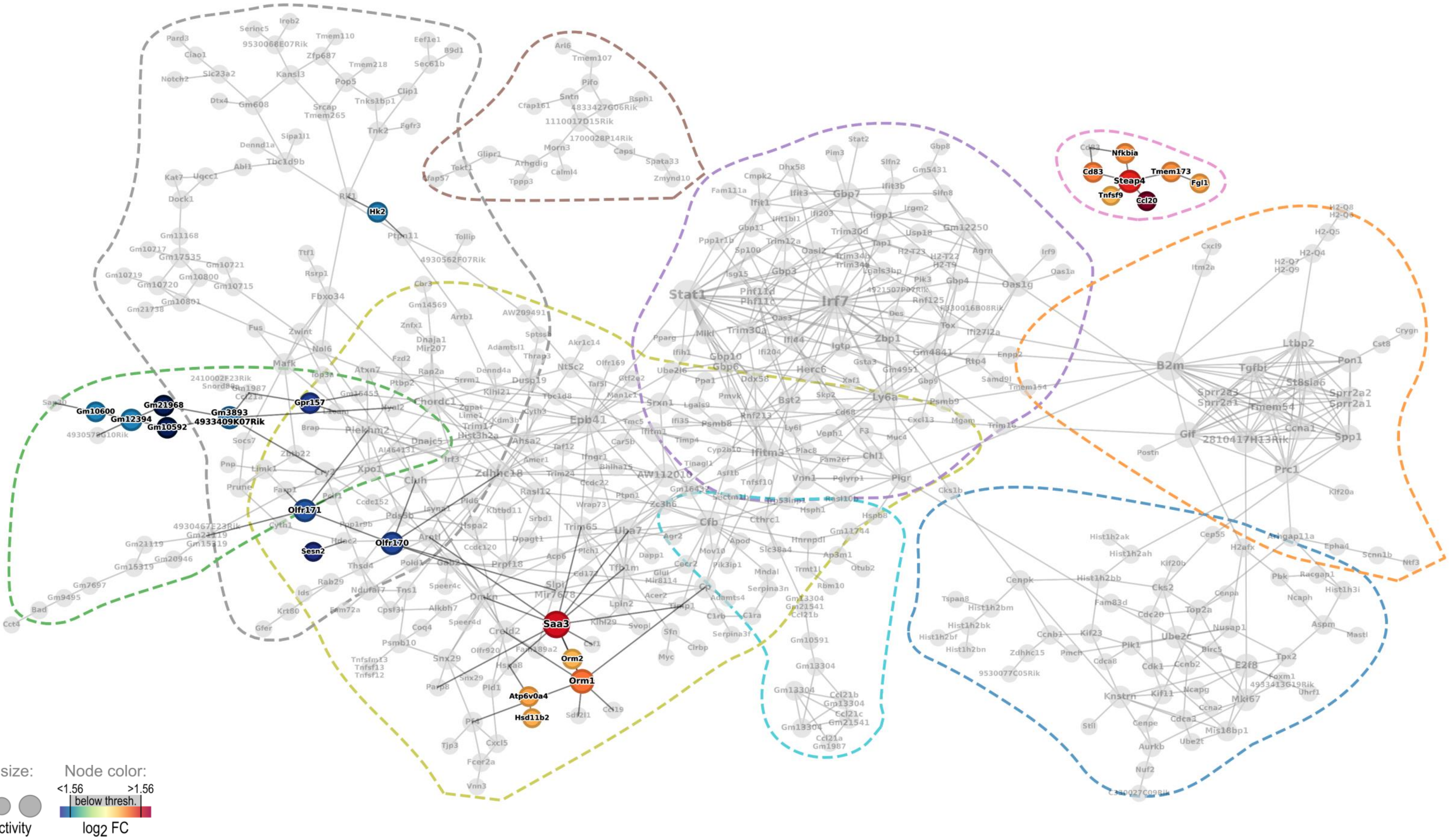

**S6: Serotype 19F 18 h vs. PBS control**

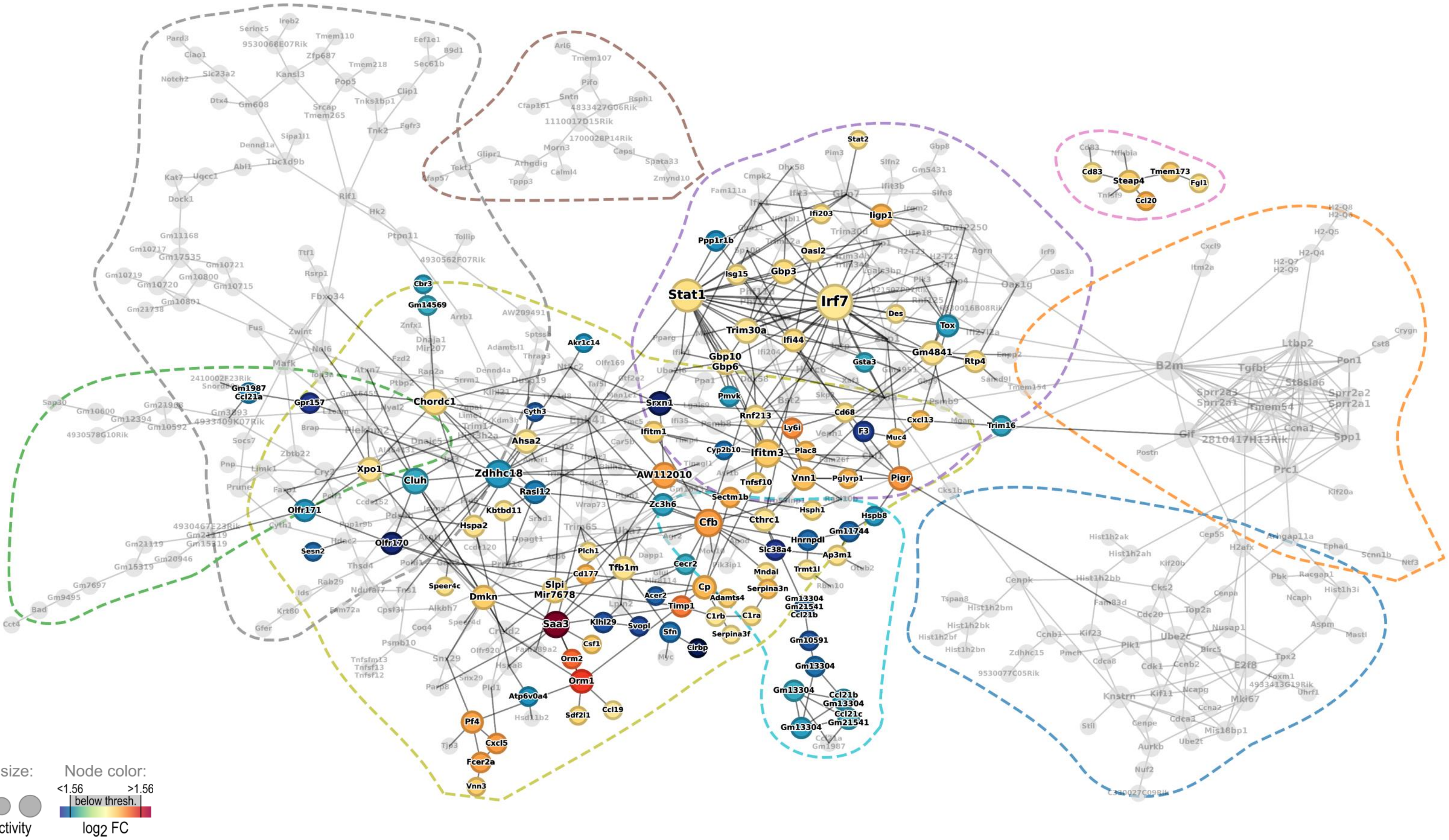

#### S7: Serotype 7F 18 h vs. PBS control

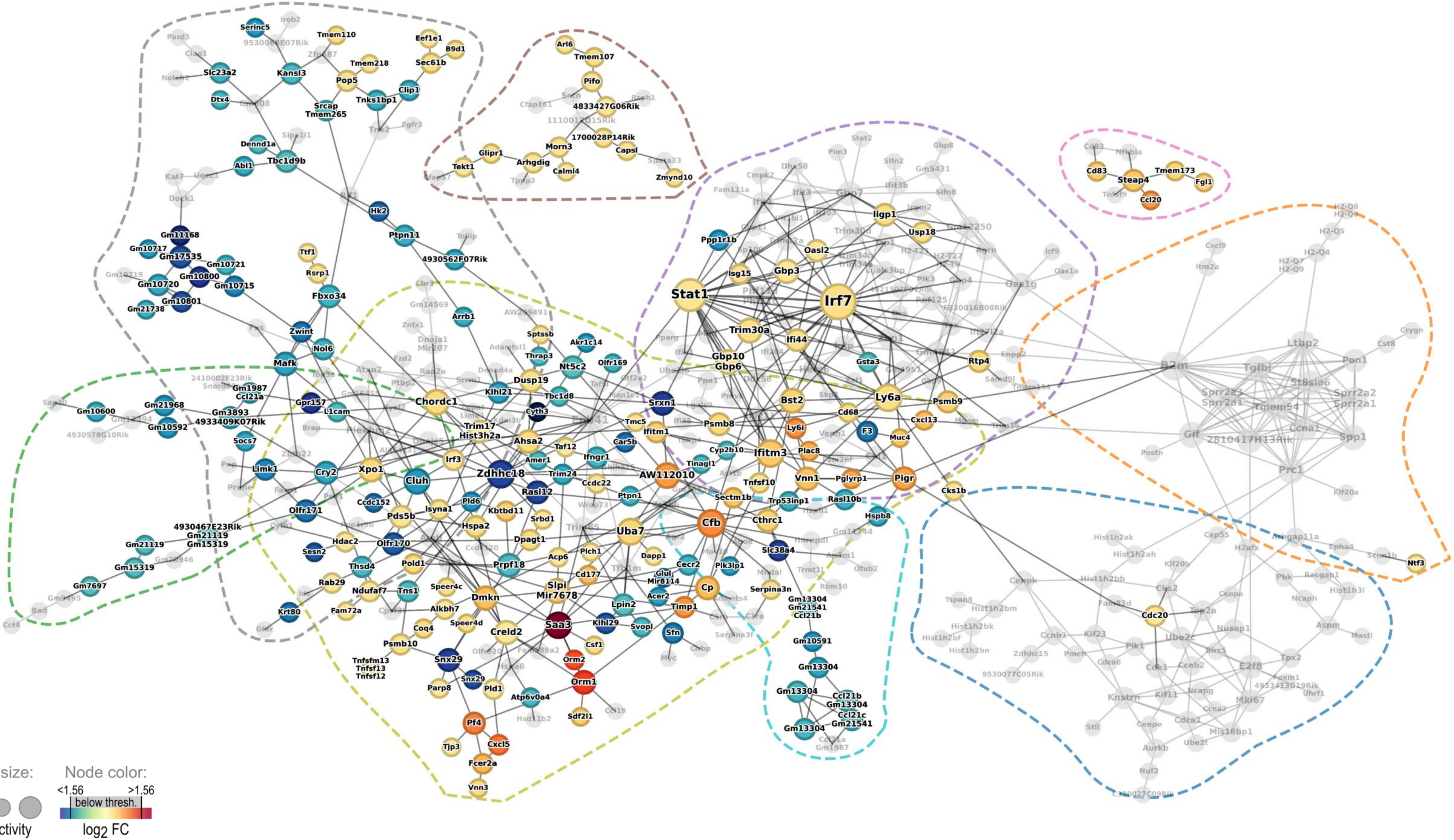

#### S8: Serotype 4 18 h vs. PBS control

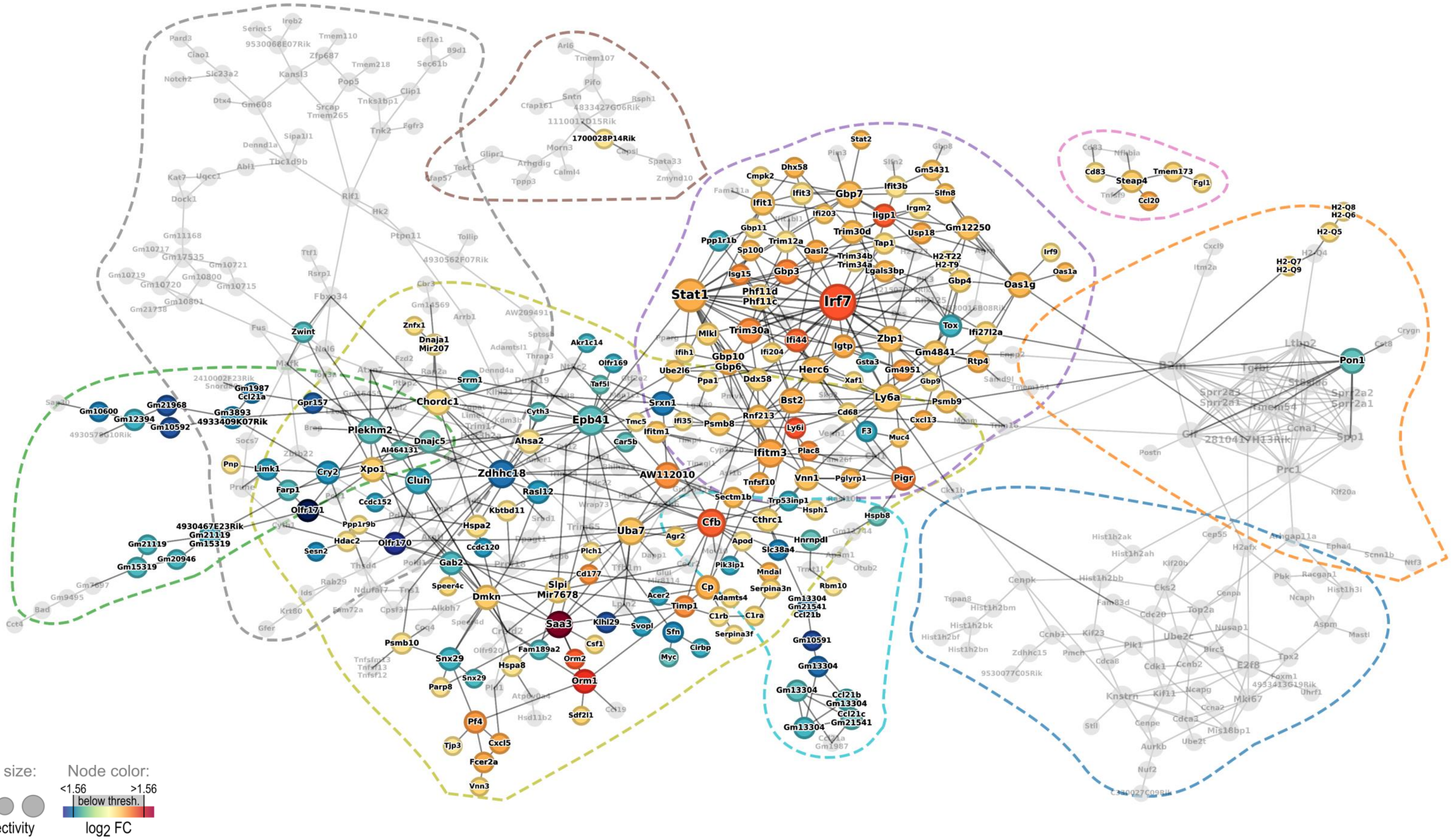

**S9: IAV day 14 vs. PBS control**

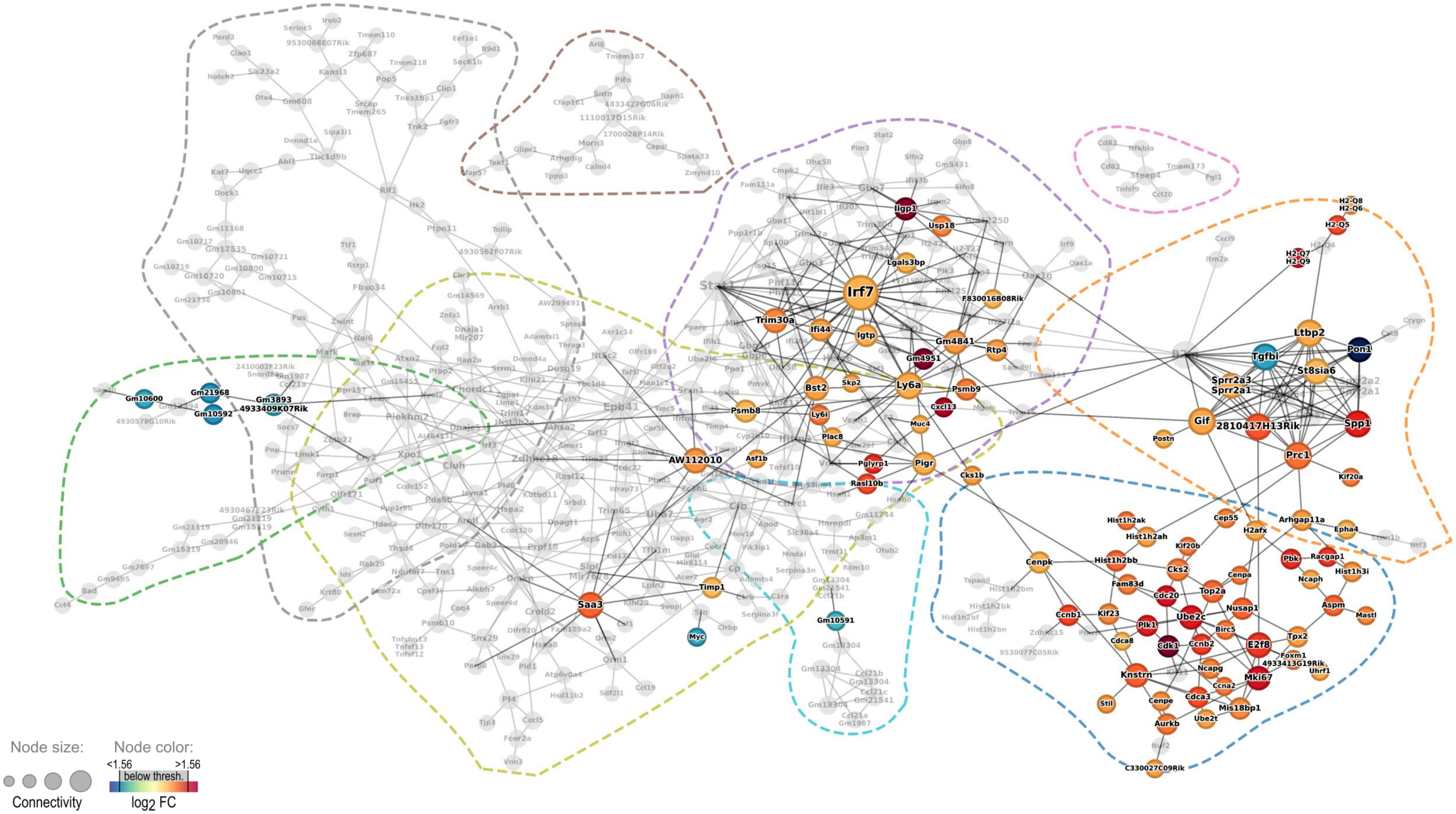

**S10: IAV day14 + Serotype 19F 4 h vs. PBS control**

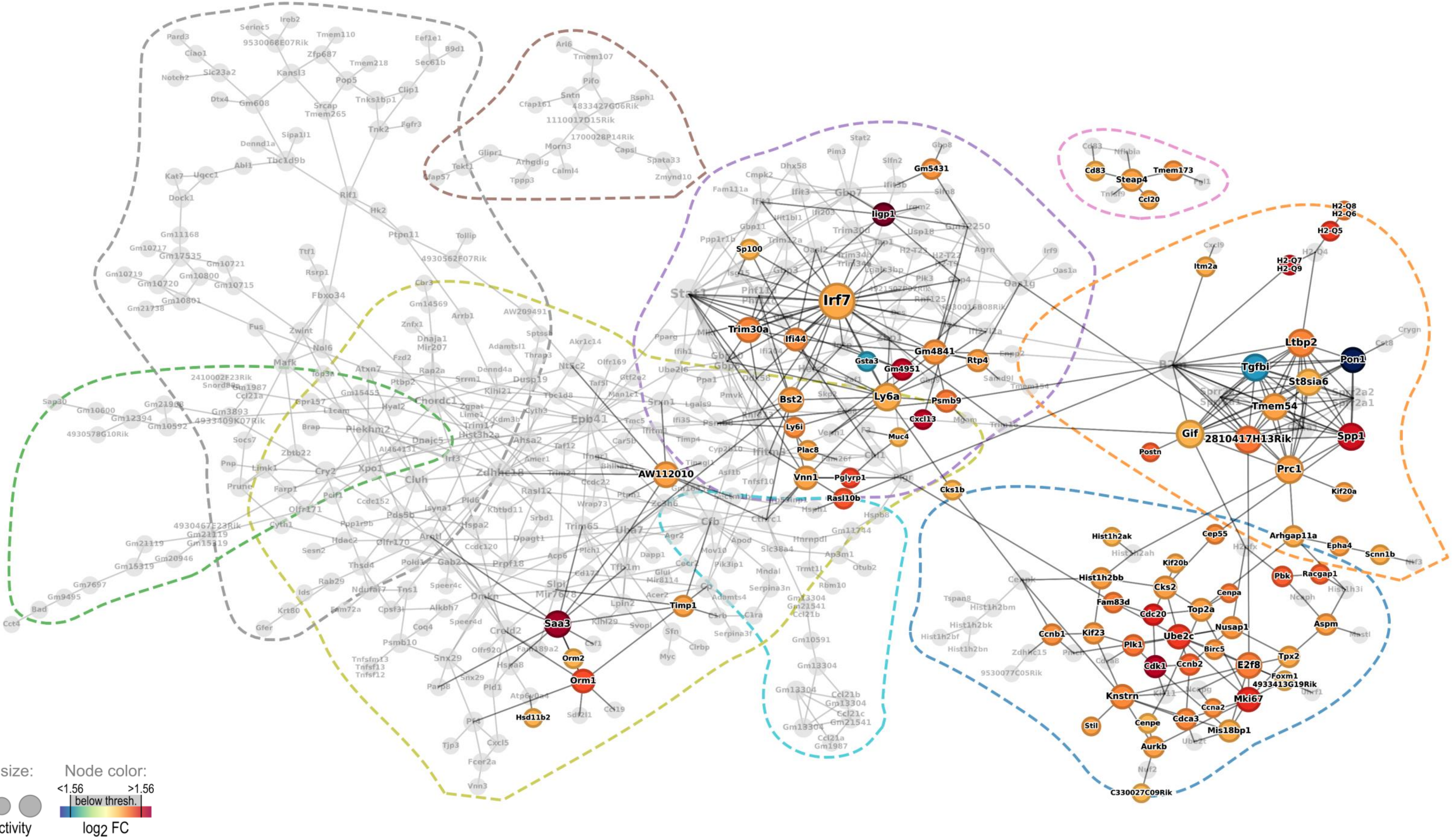

### S11: IAV day14 + Serotype 7F 4 h vs. PBS control

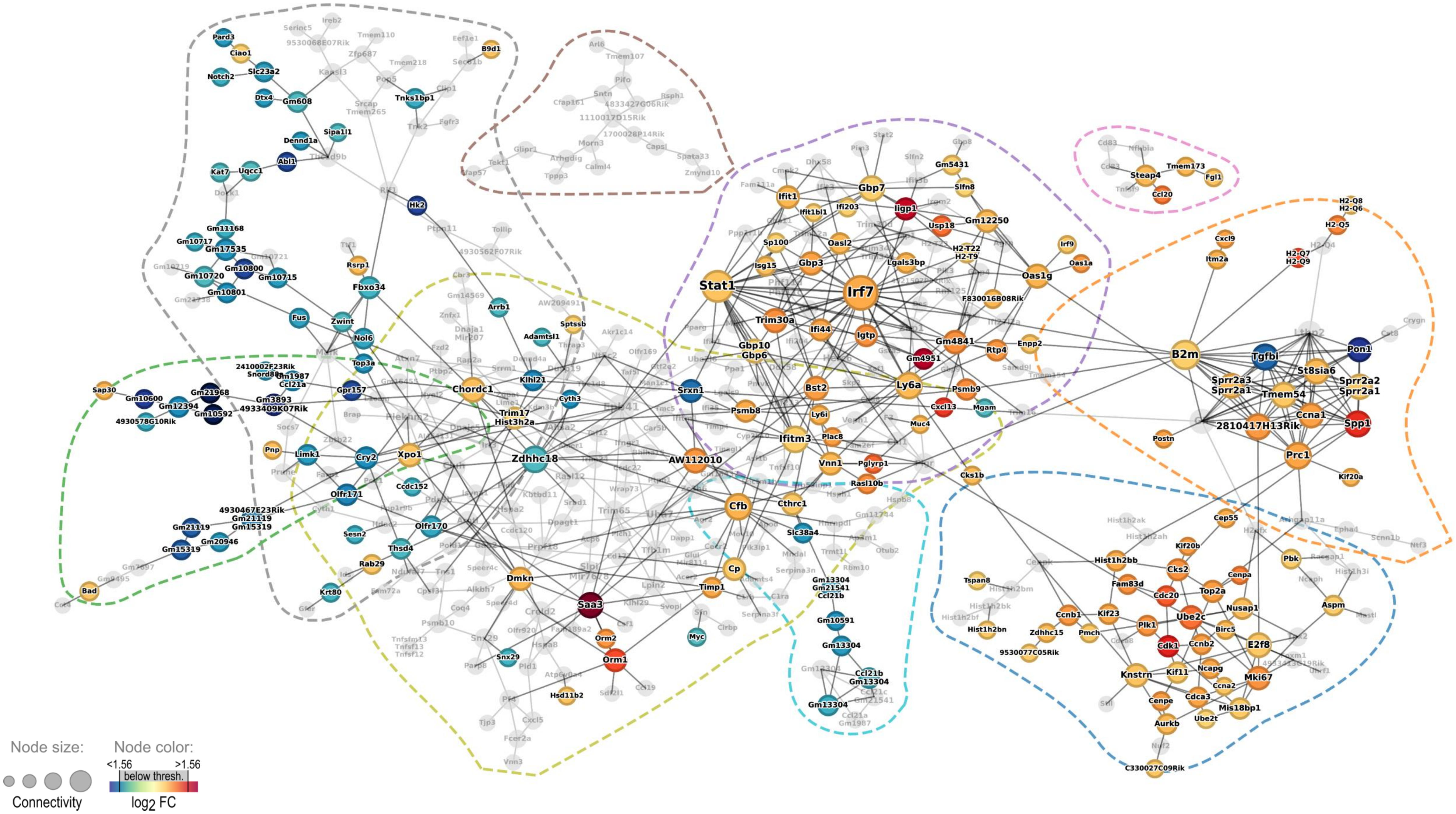

**S12: IAV day14 + Serotype 4 4 h vs. PBS control**

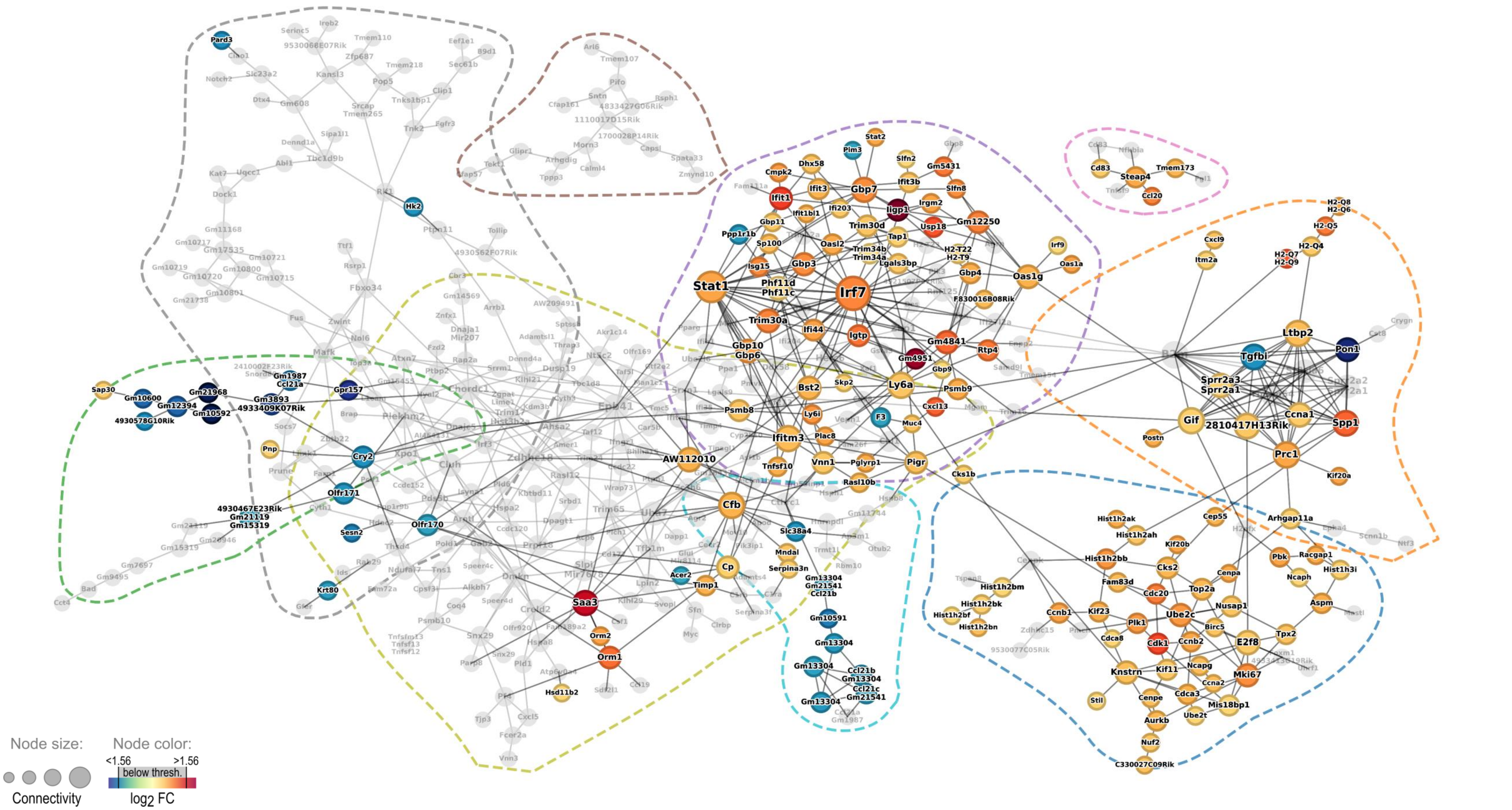

#### S13: IAV day14 + Serotype 19F 18 h vs. PBS control

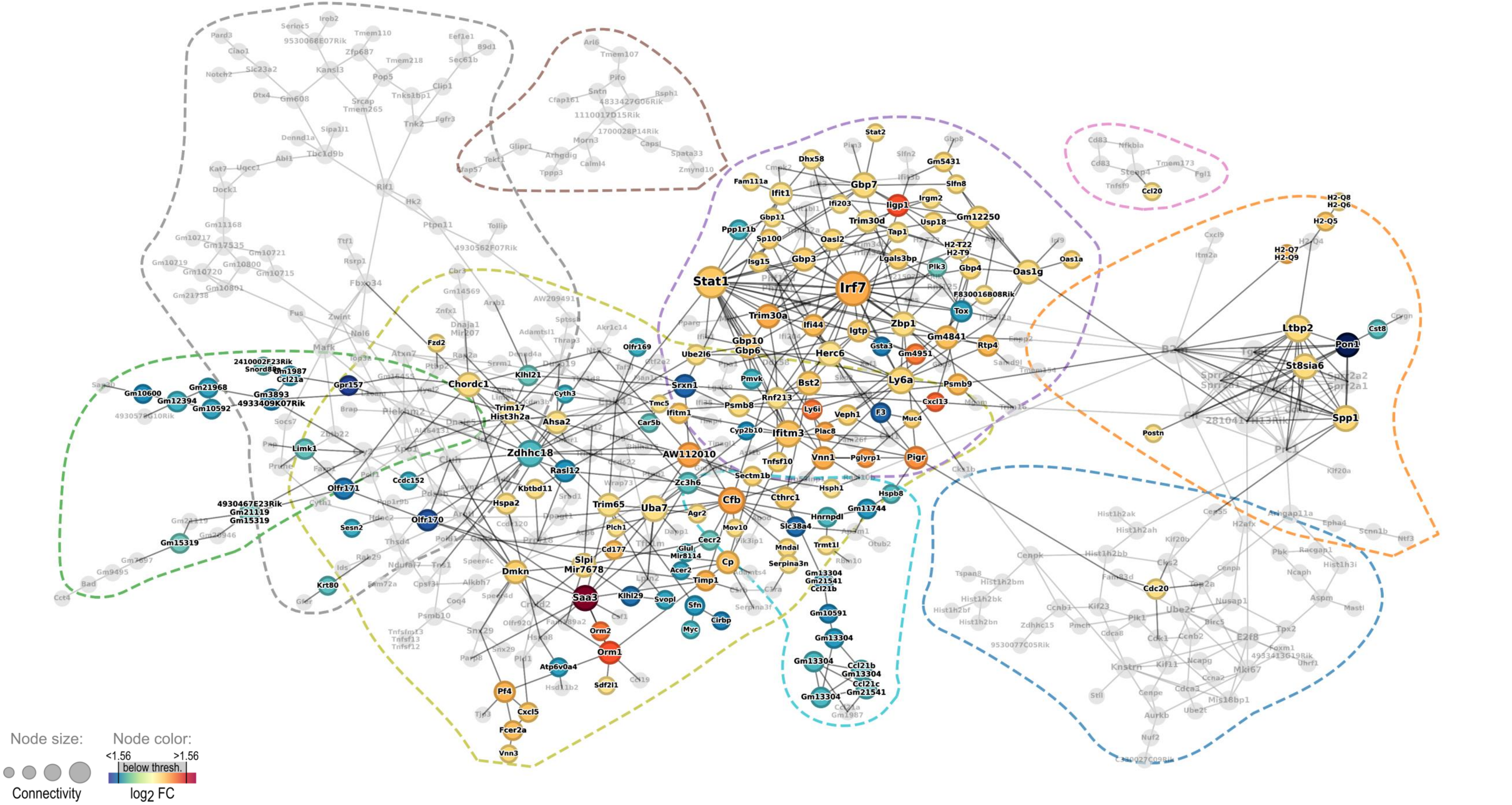

#### S14: IAV day14 + Serotype 7F 18 h vs. PBS control

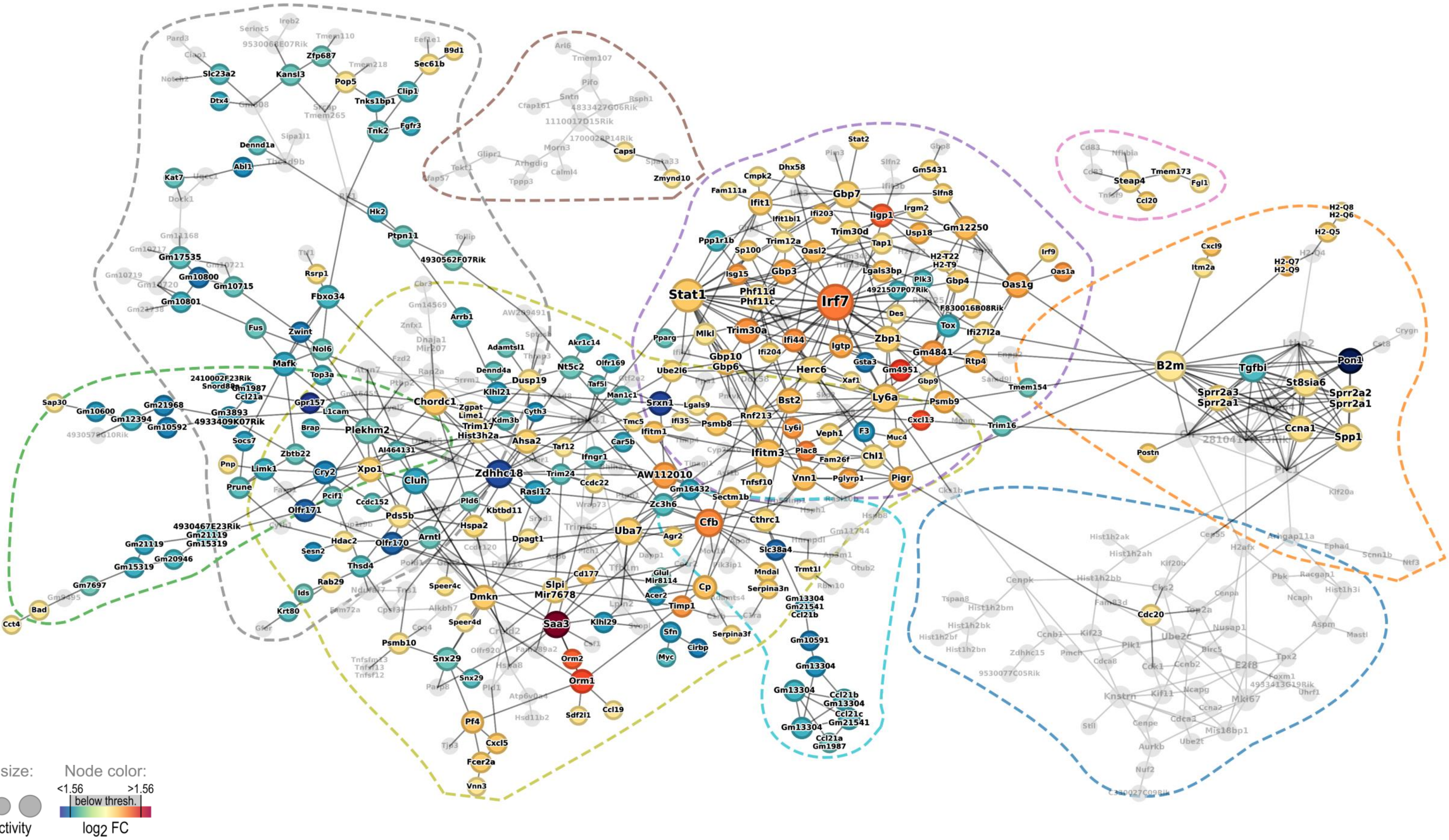

**S15: IAV day 14 + Serotype 4 18 h vs. PBS control**

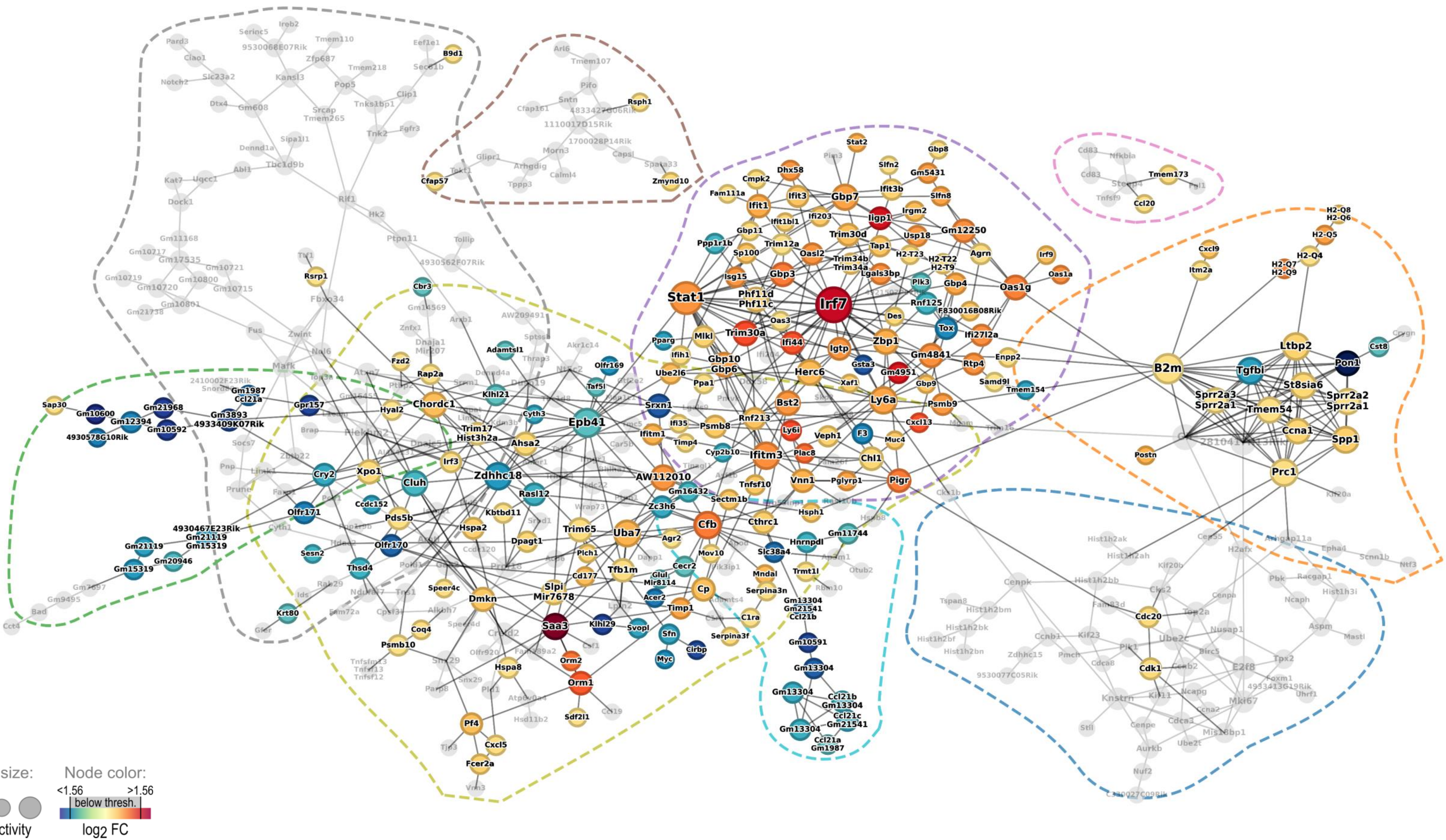

**Supplementary Figure S16: Clustering coefficients of ARACNE network nodes.** Nodes are color-coded according to their clustering coefficient. Node size indicates node connectivity. Colored dashed lines indicate outlines of network modules. Nodes are labeled with gene symbols.

#### S16: Clustering coefficients of ARACNE network nodes

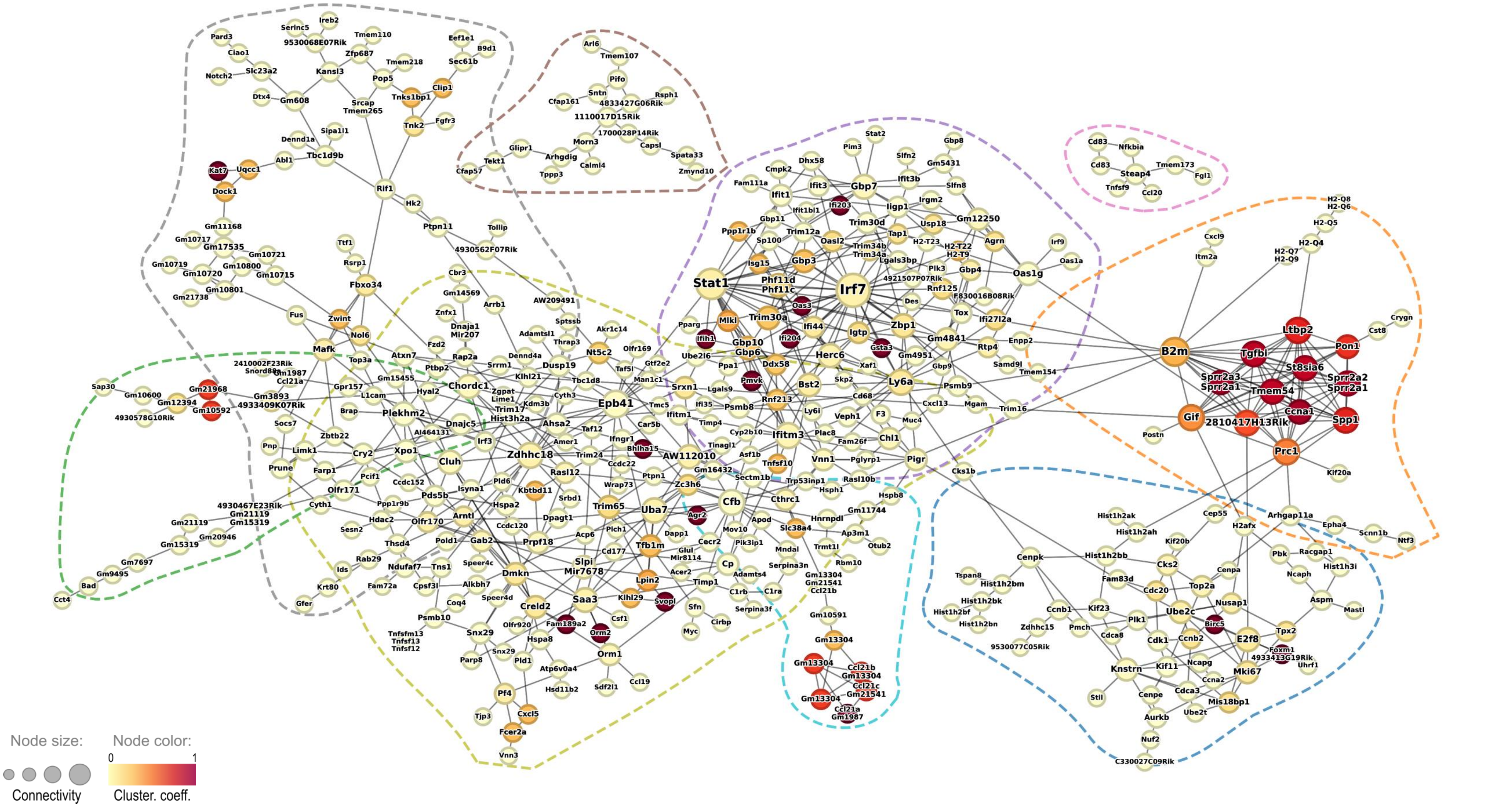

**Supplementary Figure S17: Clustered node neighbour expression heatmaps of interferon-hub genes *Irf7* and *Stat1*.** Z-scores of averaged normalized $\log_2$ signal intensities of node neighbours of *Irf7* and *Stat1* in the ARACNE network were hierarchically clustered and color coded. (+)/(-) behind genes symbols indicates positive/negative correlation of stated gene with *Irf7* or *Stat1*, respectively.

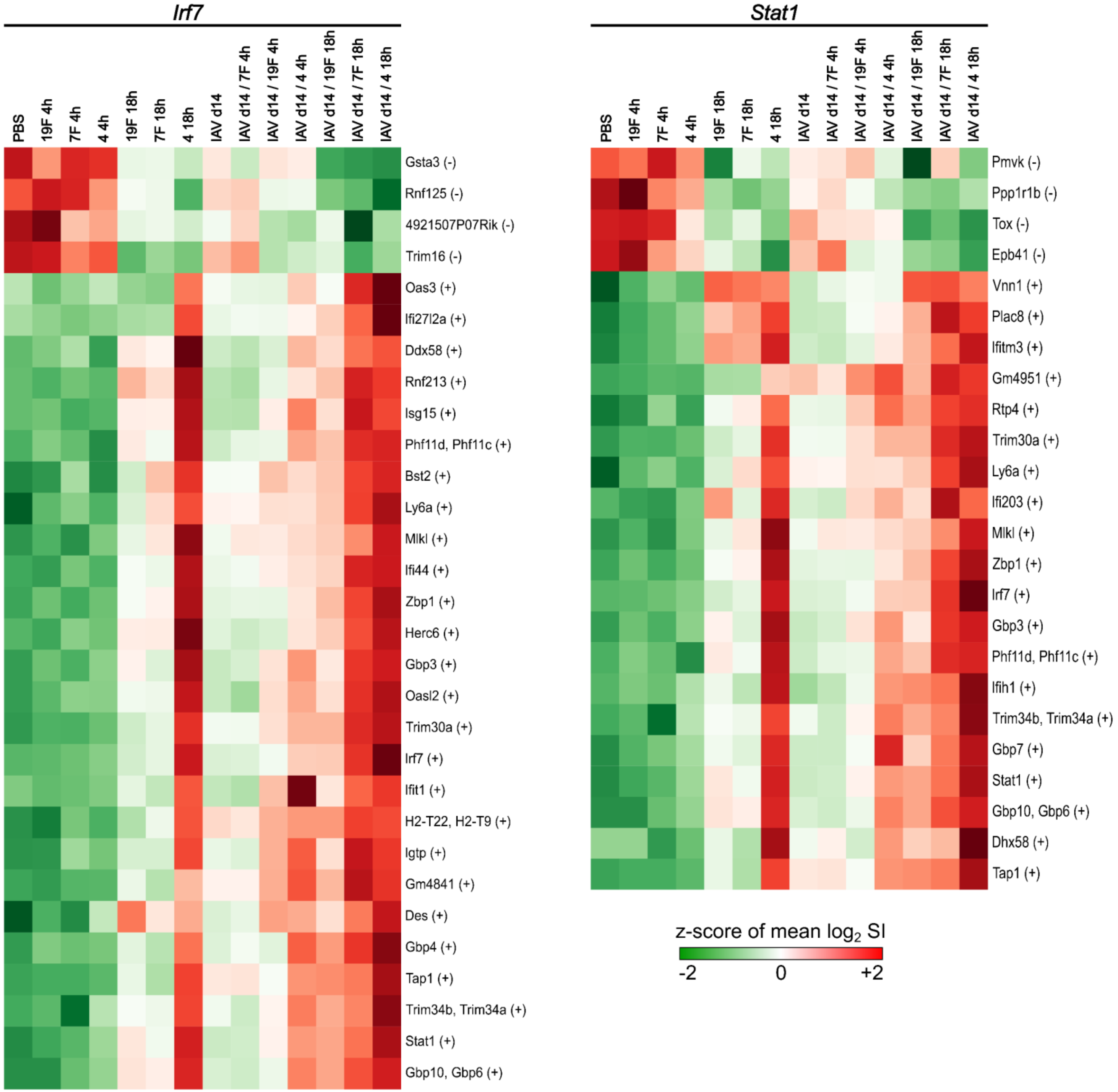

**Supplementary Figure S18: Expression of interferon genes in AECII. A)** Z-scores of normalized $log_2$ signal intensities (SI) of microarray interferon probesets. Z-scores were calculated only for shown probes per column. **B)** Average microarray SI distribution. Left: relative SI density. Right: Cumulative density. Average SI of interferon microarray probesets are indicated by black lines on the x-axis. Interferons with SI > median are labeled with gene symbols. Median, 25$^{th}$ (q = 0.25) and 75$^{th}$ (q = 0.75) are indicated by blue lines.

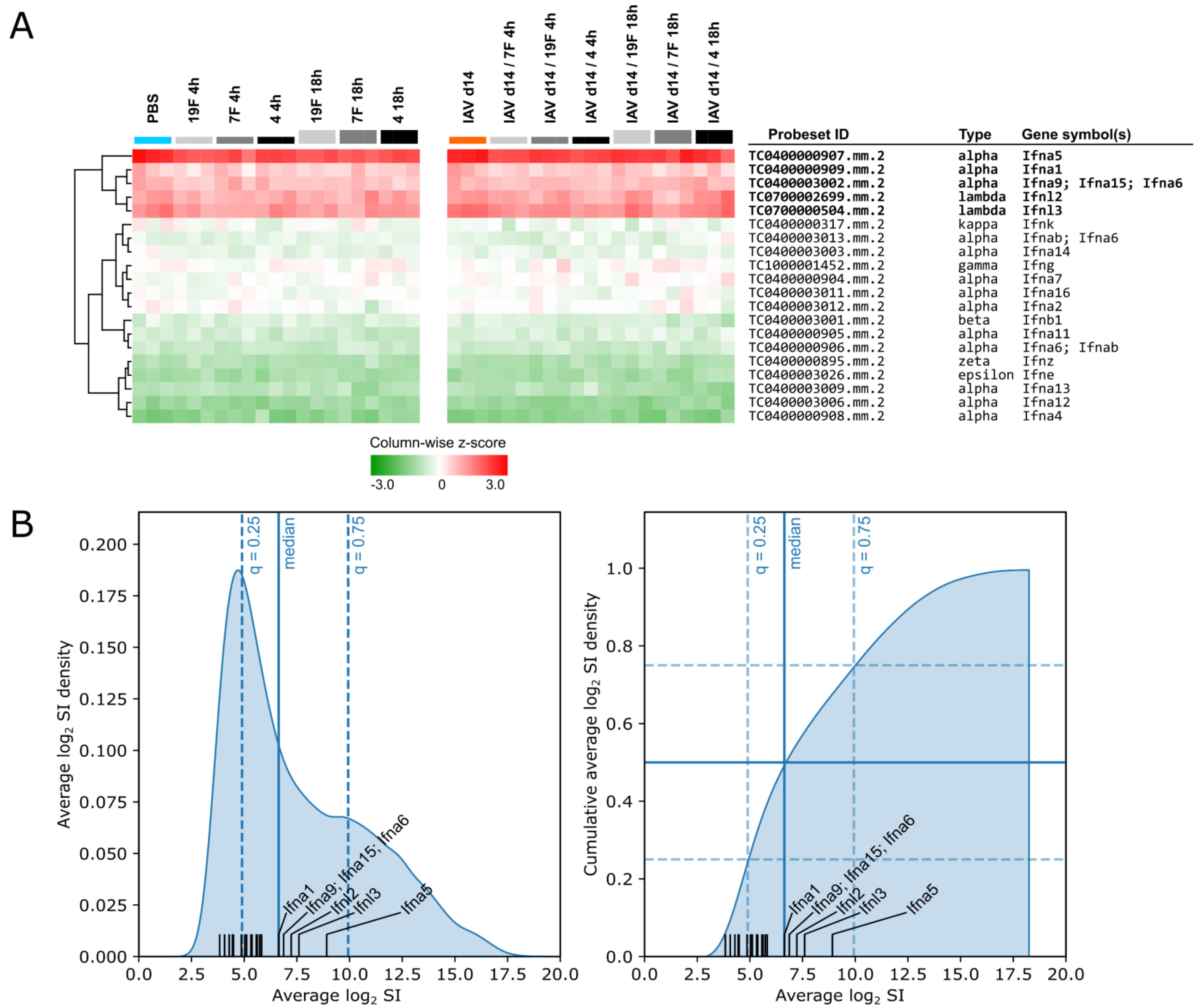
