## Supplementary Figure S19 for "Epigenetic changes and serotype-specific interferon-responses of lung epithelial cells in late post-influenza pneumococcal pneumonia"

**Supplementary Figure S19: Western blot quantification of IFN-signaling-related proteins in AECII.** Mice were intranasally infected with 7.9 $TCID_{50}$ IAV (H1N1, PR/8/34) or treated with PBS. At day 14 post IAV infection mice were sacrificed and AECII cells were sorted from n = 3 - 5 pooled lungs per sample replicate per experimental group and analyzed by western blot for expression of indicated proteins. GADPH and Vinculin were used for housekeeping gene normalization, respectively. **A-J)** Raw figures of representative western blots as shown in Fig. 7H of the main manuscript. Red squares indicate relevant target protein bands. **K-T)** Raw figures of western blots used for densitometric quantification of band intensities from 4 independent replicates as shown in Fig. 7I of the main manuscript. Red squares indicate relevant target protein bands. **U)** Table of protein band intensities derived by densitometric quantification of western blots in K-T. Protein expression is stated in arbitrary units. Row coloring corresponds to pairs of protein-of-interest and the according housekeeping protein (blue labels). Data relate to Fig. 7I of the main manuscript.

A

IRF1

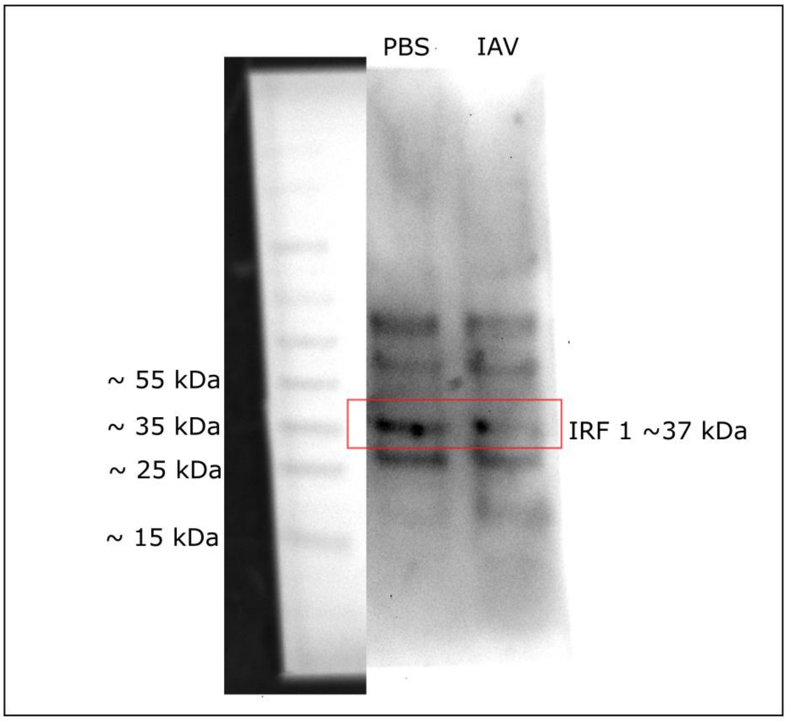


B

IRF3

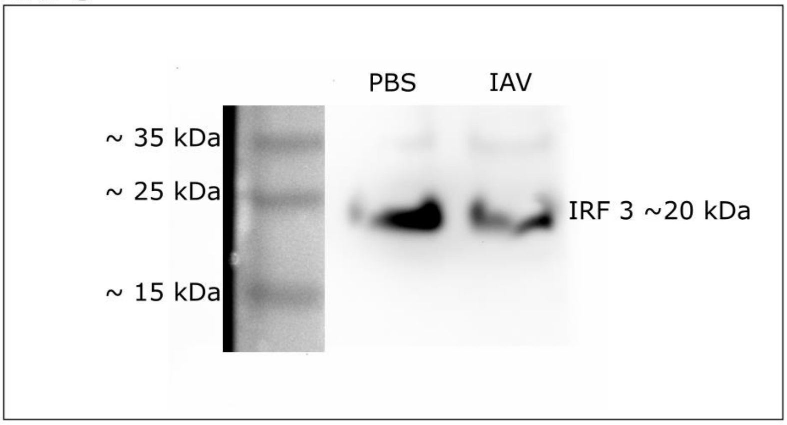


C

IRF7

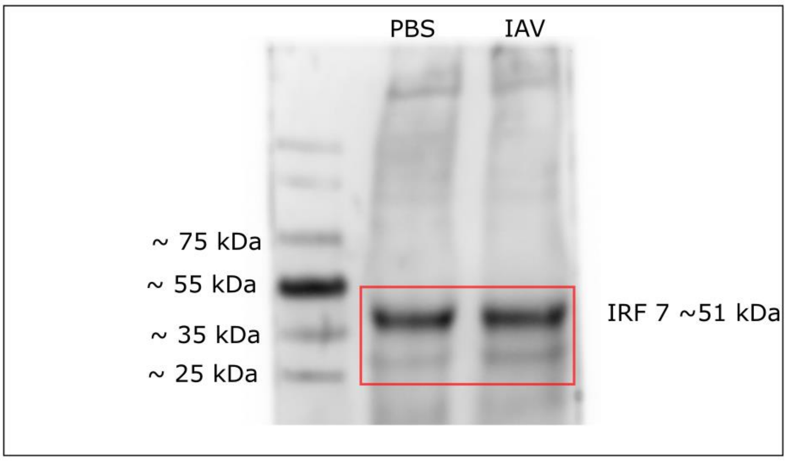

D IL10Rb

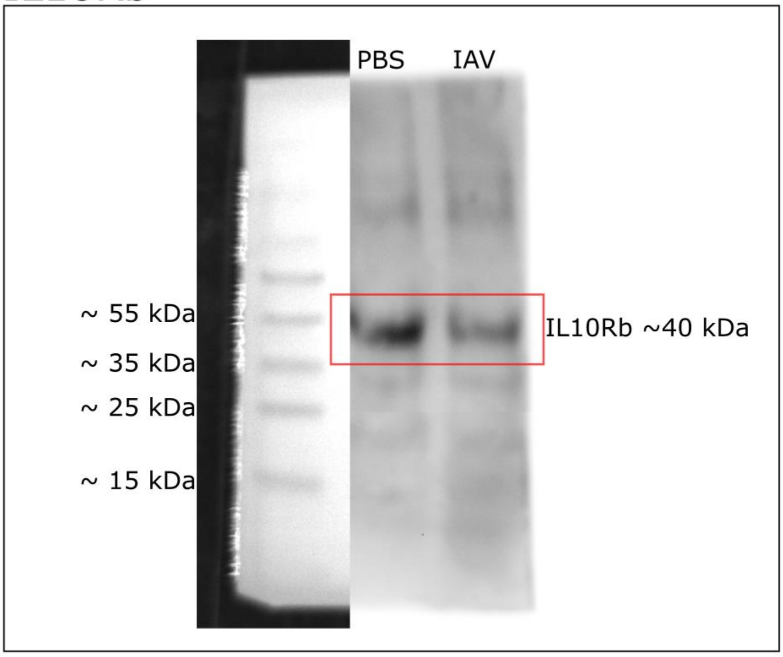


E IRF9

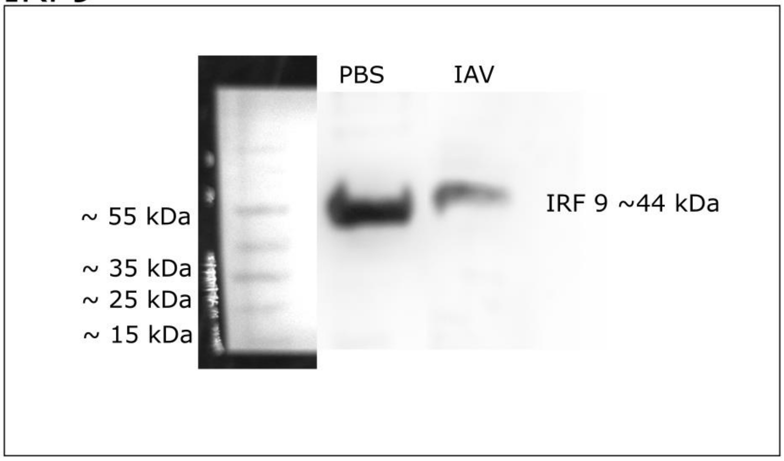


F IRFyR2

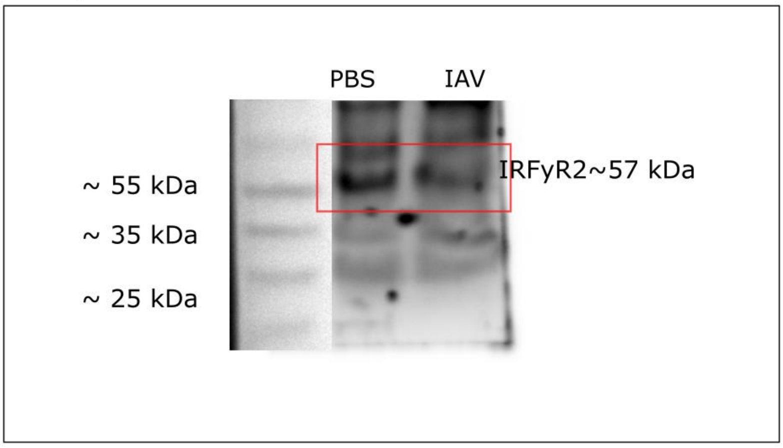

G STAT1

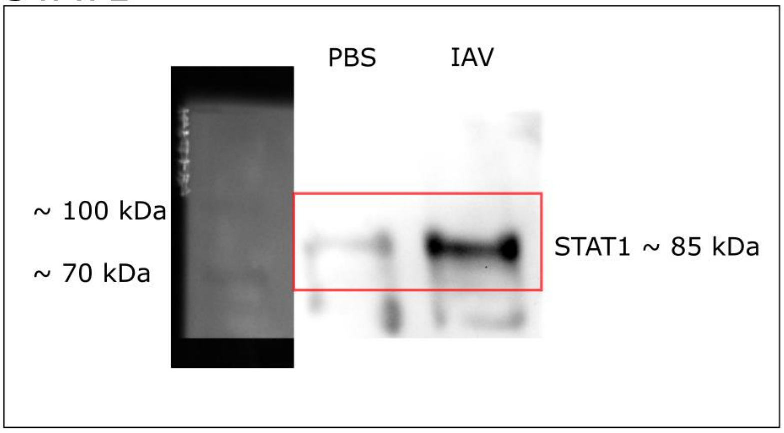


H STAT2

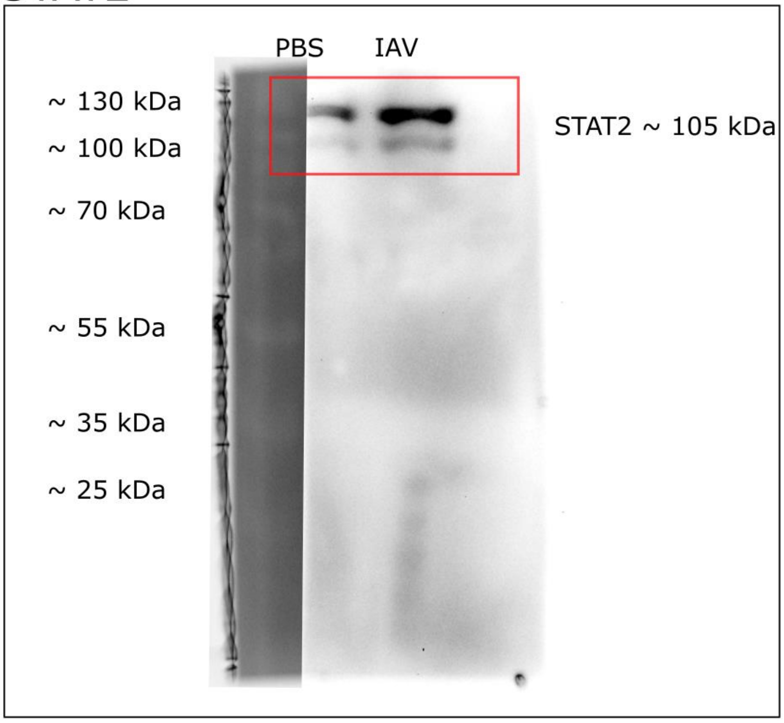


I Vinculin

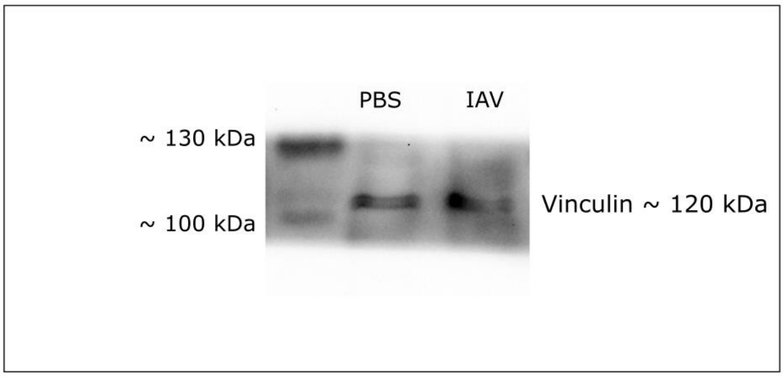

J GAPDH

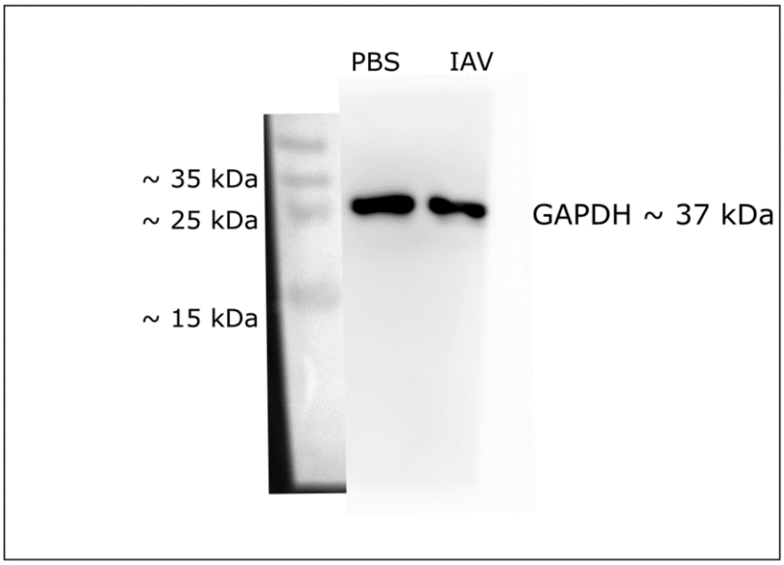

K IRF1 ~37 kDa

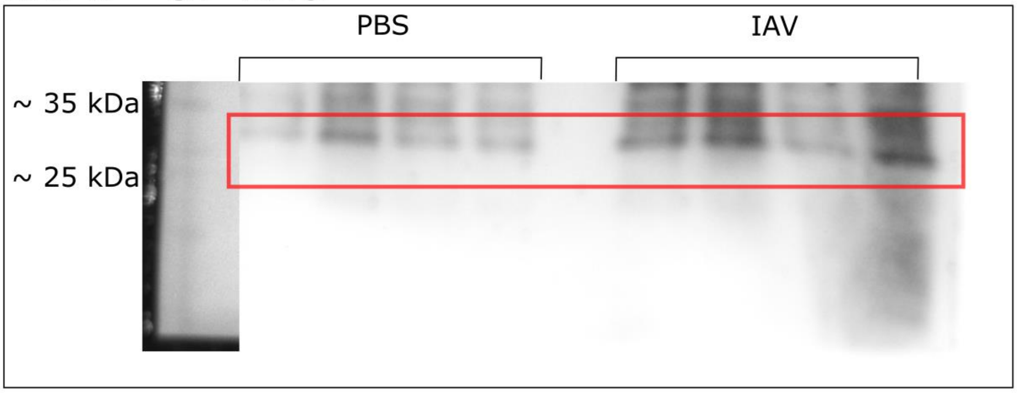


L IRF3 ~20 kDa

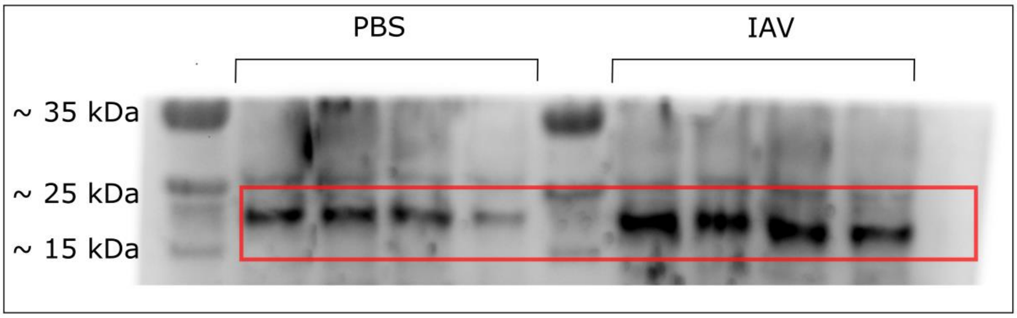


M IRF7 ~51 kDa

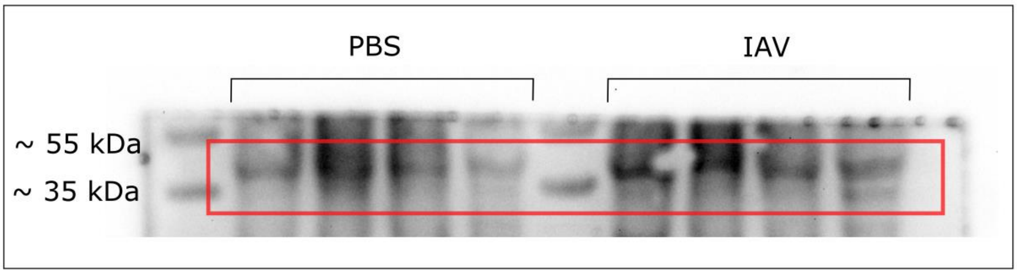


N IRF9 ~44 kDa

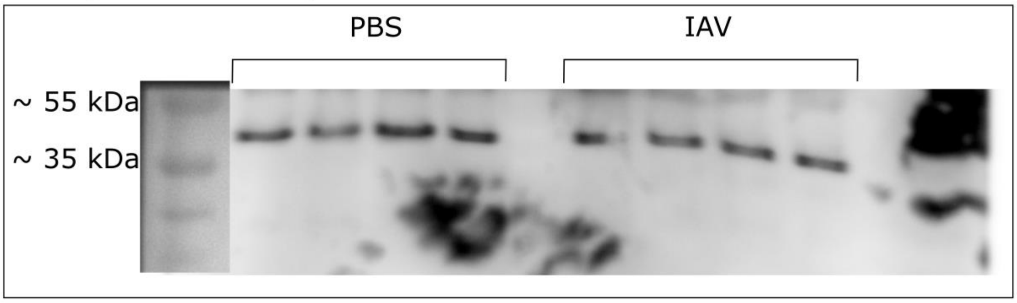

O IL-10Rb ~ 40 kDa

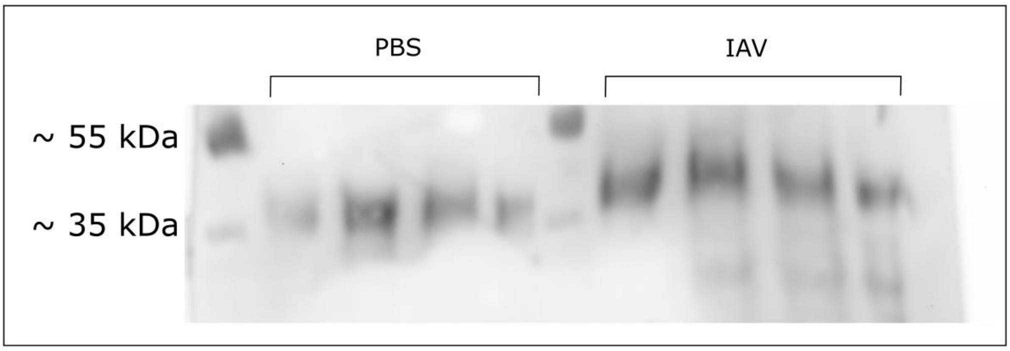


P STAT1~ 85 kDa

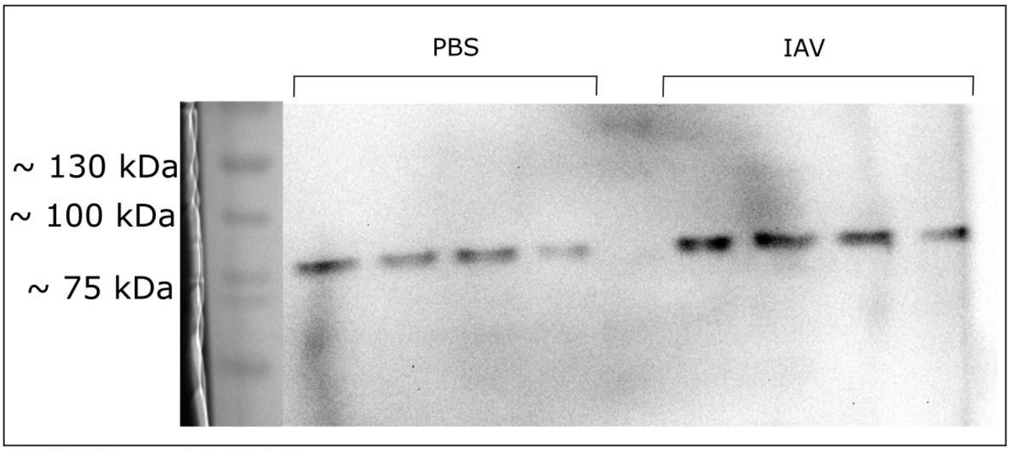


Q STAT2~ 105 kDa

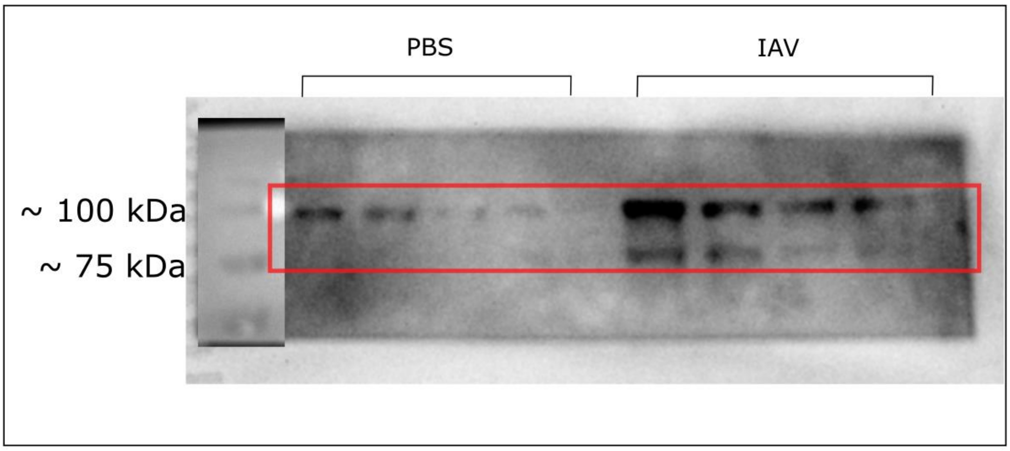

R IFNγR2 ~55 kDa

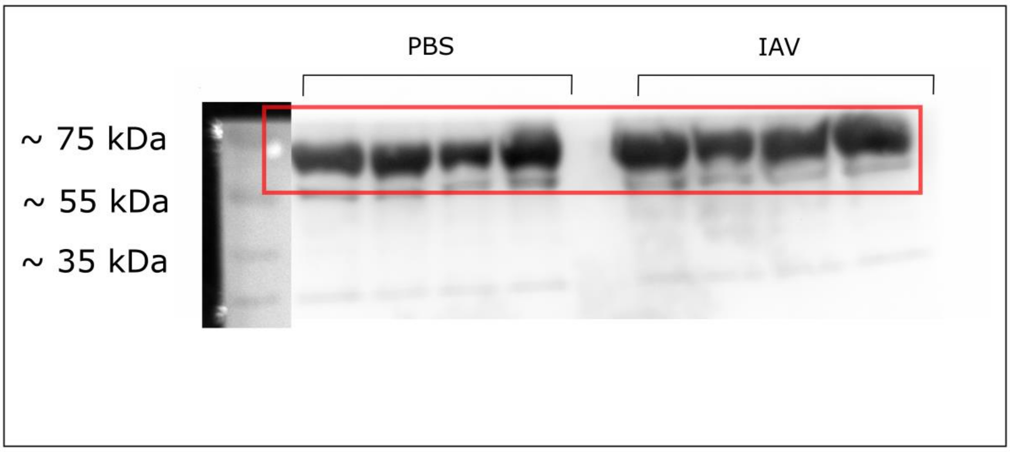


S Vinculin ~120 kDa

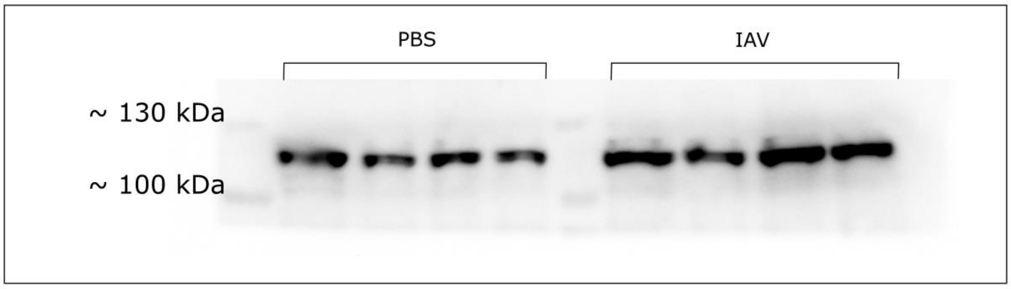


T GAPDH ~37 kDa

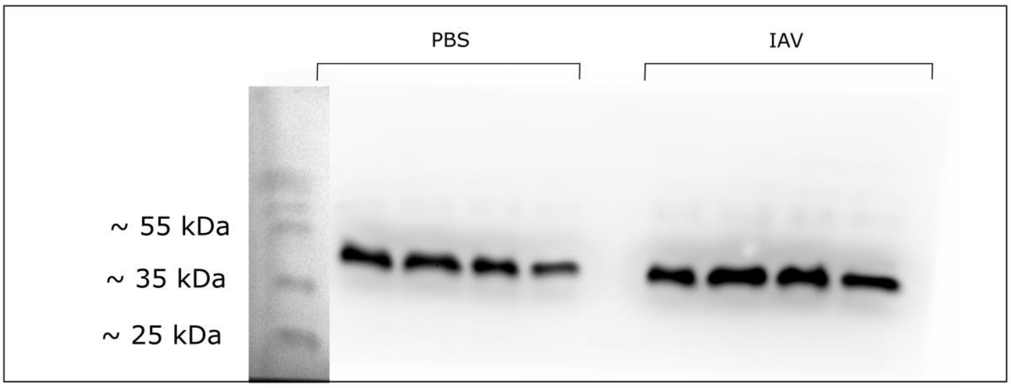

U

| Protein | PBS | | | | IAV | | | |
|---|---|---|---|---|---|---|---|---|
| | Replicate 1 | Replicate 2 | Replicate 3 | Replicate 4 | Replicate 1 | Replicate 2 | Replicate 3 | Replicate 4 |
| STAT2 | 3,051,640 | 2,680,882 | 1,022,548 | 897,598 | 10,185,388 | 6,660,075 | 4,744,731 | 2,920,397 |
| GAPDH | 15,729,602 | 14,216,095 | 12,772,510 | 9,483,317 | 14,670,803 | 19,044,045 | 16,202,217 | 13,224,610 |
| STAT1 | 3,675,317 | 4,690,832 | 5,569,196 | 1,968,347 | 8,841,782 | 4,582,803 | 7,497,782 | 6,400,104 |
| GAPDH | 6,032,832 | 7,335,024 | 6,614,660 | 4,655,832 | 4,517,125 | 2,637,539 | 5,009,974 | 6,334,217 |
| IRFγR2 | 3,740,489 | 2,229,205 | 2,319,326 | 5,431,489 | 1,652,719 | 590,991 | 897,062 | 1,851,376 |
| GAPDH | 9,498,589 | 6,863,468 | 8,239,933 | 6,228,983 | 5,951,296 | 6,344,832 | 8,026,823 | 4,922,711 |
| IRF7 | 3,490,631 | 4,092,945 | 1,590,104 | 2,163,518 | 4,443,539 | 2,076,439 | 3,228,267 | 3,879,811 |
| Vinculin | 11,684,945 | 13,278,045 | 11,915,116 | 7,811,581 | 10,690,146 | 6,548,782 | 10,202,288 | 10,601,066 |
| IRF9 | 7,616,439 | 6,028,024 | 10,226,974 | 6,881,882 | 7,616,439 | 6,028,024 | 10,226,974 | 6,881,882 |
| Vinculin | 2,817,276 | 2,655,447 | 10,139,803 | 2,582,569 | 6,421,246 | 6,596,489 | 9,692,530 | 10,223,803 |
| IL10Rb | 2,725,347 | 4,031,832 | 10,039,865 | 3,114,832 | 4,691,439 | 2,718,397 | 2,011,962 | 3,153,051 |
| Vinculin | 989,477 | 1,713,326 | 2,780,364 | 2,558,619 | 9,217,075 | 8,860,539 | 9,594,660 | 7,598,468 |
| IRF1 | 8,449,782 | 13,678,350 | 6,790,380 | 3,388,326 | 8,724,652 | 5,702,782 | 4,893,146 | 4,030,761 |
| Vinculin | 9,504,409 | 8,979,024 | 2,517,368 | 2,521,690 | 8,598,782 | 6,128,882 | 9,041,388 | 5,739,711 |
| IRF3 | 6,405,167 | 6,612,652 | 6,437,853 | 3,305,397 | 11,415,823 | 10,811,045 | 7,392,317 | 6,486,589 |
| Vinculin | 11,415,823 | 10,811,045 | 7,392,317 | 6,486,589 | 11,947,317 | 12,073,530 | 10,801,459 | 9,117,560 |
